# Invasive species-driven trophic cascades: Are cane toads indirectly contributing to small mammal collapses across tropical Australia?

**DOI:** 10.1101/616771

**Authors:** Ian James Radford, Leigh-Ann Woolley, Chris R. Dickman, Ben Corey, Dane Trembath, Richard Fairman

## Abstract

Apex predators are fundamentally important in regulating many ecosystems, and perturbations of their populations are frequently implicated in ecosystem declines or collapses. In considering small mammal declines in northern Australia, most attention has focused on interactions between a mammalian apex predator—the dingo *Canis dingo*—and a meso-predator, the feral cat—*Felis catus*. Little consideration has been given to the possible implications of changed reptilian predator assemblages resulting from invasion by a toxic anuran invader, the cane toad (*Rhinella marina*), on small mammals. We used reptile removal records from licenced reptile catchers in three widely spaced towns in the savannas of northern Australia to explore potential impacts of toads on apex and meso-predatory snakes and large lizards. In addition, simultaneous fauna survey data from one town with reptile removal records, coinciding with toad invasion, were used to identify cascading impacts through the savanna ecosystem. Intervention analyses revealed empirical linkages between toad invasion, apex predator declines, meso-predator increases and declines of small mammals and other prey groups. Based on the timing and strength of intervention we postulate a novel conceptual model linking recent mammal declines with trophic cascades following toad invasion, where the loss of large, anurophagous (toad-eating) reptilian apex predators allowed increases in mammal-eating meso-predatory snakes. The conceptual model is discussed in relation to prevailing hypotheses regarding northern Australia’s dramatic small mammal declines. Future studies will need to quantify these putative interactions and test their comparative importance so that appropriate management can be implemented to stem the ongoing losses of mammal fauna.

## INTRODUCTION

> “We were the Leopards, the Lions; those who’ll take our place will be little jackals, hyena - and sheep”, Giuseppe Tomasi di Lampedusa (The Leopard)

The importance of apex predators and their interactions with smaller meso-predators in maintaining global ecosystems has received increasing attention in recent decades. Meso-predator suppression by apex predators is widespread both geographically and taxonomically [1]. Apex predators clearly play a crucial role in maintaining many ecosystems [1], but debate continues about how pervasive their effects are [2-3]. In Australia, for example, several studies suggest that a mammalian apex predator, the dingo *Canis dingo*, suppresses feral cat *Felis catus* activity and the impacts of the latter on native species [4-8], but other work contests this [3]. The influence of apex predators may change depending on the number of different sized meso-predators within predator hierarchies, leading to different outcomes for prey populations [9]. One important lesson from classic predatory studies (e.g. wolves, moose and bears in North America) [10] is that all relevant predators must be considered to achieve a full mechanistic understanding of prey dynamics.

Meso-predatory interactions have been reported most commonly among mammalian predators, raptors, and in marine systems [1]. In recent years, however, meso-predator release within reptilian predator assemblages has been identified in northern Australian savanna ecosystems resulting from invasion by a toxic anuran, the cane toad *Rhinella marina* [11-15]. Here, losses of large reptilian apex predators – varanid species – due to post-ingestion poisoning by toads, have resulted in measurable increases among smaller, meso-predatory reptiles which presumably had been eaten by varanids [11-13]. Despite the rarity of comparable examples [but see 16-17], it is not surprising that this case of reptilian meso-predator release was identified in Australia given the relative abundance of large reptiles in Australian ecosystems, as well as the paucity of large mammalian predators [18]. However, it remains unclear whether meso-predator release among reptiles might have wider implications within the tropical savanna [14, 19].

Recent rapid and dramatic collapses of small to medium-sized mammals in northern Australia [20-21] continue to puzzle ecologists, although local declines here have been foreshadowed for several decades [22-26]. Recent thinking concerning these declines implicates feral cats interacting with changed fire regimes, large herbivores, and possibly ecosystem condition/ productivity, to negatively affect small mammals [4, 25, 20, 27-30, 21, 31]. Cats (along with foxes *Vulpes vulpes* in sub-tropical regions) are thought to have been the primary drivers of historical extinction events across Australia, particularly in arid regions where arrival of cats often coincided with sudden mammal declines even before European settler arrival or the operation of other threatening processes [32, 4]. However, there is little direct evidence to link recent accelerated collapses of northern Australian mammal assemblages in the first decade of the twenty-first century to cat predation. Cats coexisted with susceptible mammals here for a century [33] prior to the recent collapses [34, 20] and there is no evidence that recent changes in cat populations, fire or grazing regimes coincide with these sudden declines. Experimental studies have shown that cat predation can cause local extinctions of rodents inside fenced savanna areas [27, 35], and convergence of cats at recently burnt areas [28, 30] can dramatically increase predation-related mortality of small mammals [31]. There is also a link between more severe fire regimes and lower mammal abundance/richness [34, 36-37]. However, these studies do not establish that cat and fire interactions have caused mammal collapses at regional scales [38]. There is no evidence currently available showing that cat populations or predation pressure have increased at the same time that mammal declines were occurring. There are also no data to link any sudden exacerbation of fire/grazing related disturbance regimes at regional levels with recent observed mammal declines. Thus, there is no empirical evidence to directly link recent mammal collapses in the Northern Territory [34, 20-21] with changes in the operation of any known threatening processes (e.g. cats, fire regimes, large herbivores) at the same time that the observed mammal declines occurred.

One factor acknowledged as potentially influencing mammal assemblages, but for which little evidence has been adduced, is invasion by the cane toad [20-21]. Clear evidence links recent declines of one mammal species, the northern quoll, *Dasyurus hallucatus*, to the arrival of the cane toad [39-40]. Quolls actively hunt and ingest toads and are subject to high rates of mortality due to lethal poisoning [40-41]. However, declines among other small mammals have not been linked to toad arrival. This is because most mammals do not eat cane toads, or escape poisoning by avoiding the toxic glands [40]. Similarly, small mammals are not known to be eaten by cane toads [39].

In this paper we present a new hypothesis explaining recent north Australian small mammal declines based on observed changes seen in savanna fauna following invasion by cane toads at three separate locations across north-western Australia. The hypothesis is that recent mammal collapses, since 2005 in the Northern Territory, and since 2010 in Western Australia, may be attributable to the arrival of cane toads via a series of cascading impacts on reptilian predator assemblages including mammal-eating species. Changes following cane toad invasion included an immediate decline of large-gaped, large-bodied, generalist (partially anurophagous and reptile-eating) elapid snakes such as the king brown snake (*Pseudechis australis*) and varanid lizards (from here referred to as apex predators) as seen during previous studies [15, 39, 42]. We interpret these collapses as due to poisoning upon ingestion of toxic cane toads [39, 15]. Another change at study sites was an immediate increase among smaller-gaped, smaller-bodied (i.e. meso-predatory), dietary specialist snakes, lizards and anurans, including mammal-eating pythonid and cobubrid snakes. Increases among meso-predatory reptiles was also previously reported in several studies [11-13, 19]. We interpret increases as due to a meso-predator release following loss of large generalist apex reptilian predators [15]. Additional changes associated with cane toad invasion at one of our study sites where we had continuous fauna monitoring data were declines among fauna which is prey to meso-predatory reptiles. Prey included small mammals, very small skinks (< 8 cm long) and many invertebrates. Declines among invertebrates following toad invasion has been reported before [43], though this was interpreted as due to an increase in predator biomass due to cane toad presence, not due to general increases among a range of meso-predators. We interpret declines among prey groups as being driven by increases among their meso-predators, including mammal- and lizard-eating snakes, medium-sized lizards and frogs, as these species all increased following cane toad invasion. We synthesise these empirical observations in the form of a conceptual model that articulates the trophic links between small mammals, cane toads and reptilian predators (Fig. 1). This provides us with a complementary hypothesis to the cat – fire/disturbance driven hypothesis that has dominated the literature on north Australian mammal declines over the last decade [4, 27, 36, 21, 30].

**Fig. 1.**
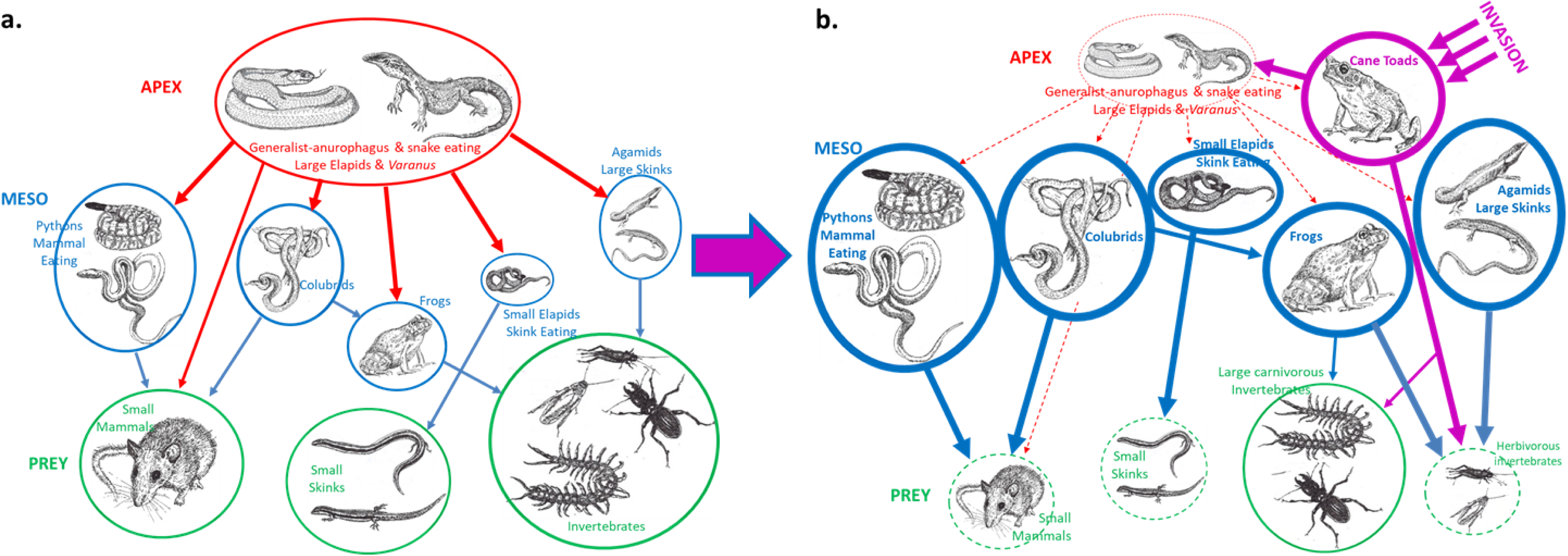
A conceptual model predicting changes in trophic interactions among terrestrial savanna fauna with the arrival of cane toads (*Rhinella marina*) in savannas adjacent to Kununurra. Savanna apex reptilian predators include large elapid snakes (e.g. *Pseudechis australis*) and varanid lizards (*Varanus gouldii*). Meso-predatory reptiles include mammal-eating snakes such as pythons (e.g. *Aspidites melanocephalus* and *Liasis olivaceus*), colubrid snakes (e.g. *Dendrelaphus punctulatus*) and small skink-eating elapids (e.g. *Furina ornata*). Other savanna meso-predators include frogs (e.g. *Platyplectrum ornatum*), large scincids (e.g. *Ctenotus robustus*) and agamids (*Lophognathus gilberti*). Arrow thickness represents the strength of the interactions between trophic levels (apex and meso-predators and prey species). Thin, dashed lines or arrows indicate putative declines or weakened interactions. Violet arrows/lines represent interactions with the invasive cane toad, red lines/arrows represent apex predators and their interactions, blue lines/arrows meso-predators and their interactions, and green lines are key savanna prey species/groups. **a)** Represents a conceptual model of trophic interactions in savanna ecosystems prior to cane toad invasion. Pre-invasion reptilian and amphibian assemblages were dominated by the apex predators, which were the large-gaped anurophagous/generalist reptiles, which suppressed many of the meso-predatory savanna species, including reptilian, amphibian and mammal species. In this pre-invasion ecosystem, prey groups including small mammals, small skinks and invertebrates persisted at moderate abundance. **b)** Shows how these interactions are predicted to alter following cane toad invasion. With the loss of ca. 80% of the large, anurophagous/generalist apex reptilian predators, meso-predatory snakes, frogs, skinks and agamids increased by ca. 250 % and cane toads were introduced as an additional meso-predator. Under this scenario, there was increased predation pressure on prey groups including small mammals, small skinks and some invertebrates (herbivorous) which resulted in declines in these groups of ca. 30-80%. Note that large predatory invertebrates including carabid beetles and centipedes neither declined nor increased following cane toad invasion.

## METHODS

### Study areas

The main study location was the town of Kununurra (2016 census population 5,300) and its surrounding savanna landscapes including Mirima National Park in far north-eastern Western Australia (Fig. 2a, b). The region has a tropical monsoonal climate, with high temperatures year-round (daily mean maximum 29.6-36.0 ^o^C), and rainfall (913 mm annually) occurring predominantly from November to April. Several tropical savanna habitats occur around Kununurra. Aside from urban and agricultural (broad-acre cropping) habitats, these include black soil plains, eucalypt woodlands dominated by tussock grasses, pindan (*Acacia tumida*) savanna woodlands dominated by *Triodia* hummock grasses and annual *Sorghum* on sandplain, and shrub/*Triodia* spp. dominated woodland on rocky sandstone. Kununurra is adjacent to perennial riparian habitats and permanent water due to the damming of the Ord River (Fig. 2a, b). Minor study locations at Katherine (popn. 6,300) and Darwin (popn. 136,800) in the Northern Territory have similar tropical monsoonal climates to Kununurra (Fig. 2a), with daily mean maximum temperatures of 30.1–37.7 ^o^C (Katherine) and 30.6–33.3 ^o^C (Darwin), and mean annual rainfalls of 1023 mm and 1729 mm respectively. Both towns, like Kununurra, are small and are predominantly surrounded and interspersed by tropical savanna habitats.

**Fig. 2.**
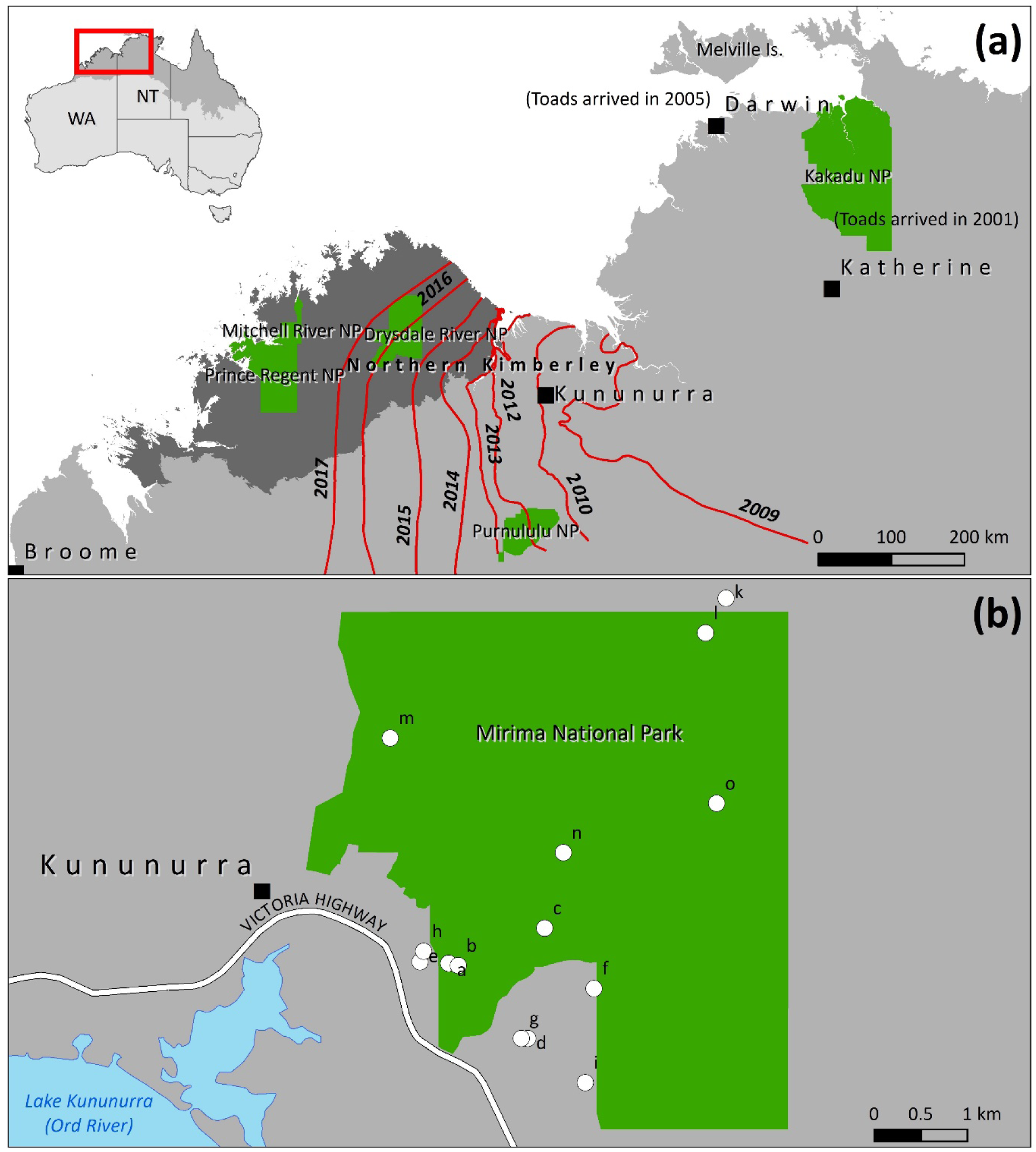
a) North Western Australia including the minor study areas of Katherine and Darwin and b) the main study area of Kununurra and Mirima National Park survey sites (white dots) in the Kimberley region of Western Australia (WA). The red inset (a) shows the study region within the broader savanna biome (darker grey) in Australia. The map also shows sequential invasion of cane toads (*Rhinella marina*) across northern Australia. Cane toads began arriving in Katherine and Kakadu National Park in the Northern Territory (NT) in 2001 and were in Darwin by 2005. Red contour lines mark the estimated cane toad invasion front annually from the end of 2009, when toads first entered WA from the NT, in 2012 when toads arrived in Purnululu National Park, through to 2016 and 2017 when toads first arrived in the Drysdale River National Park in the North Kimberley bioregion (dark grey). The Cane Toad Strategy for WA 2009-2019 provided annual spatial data on invasion fronts. National Parks are shown in green.

### Reptile removal records

Removal records of reptiles were obtained from Kununurra, Katherine and Darwin. Trained personnel (government officers or volunteers) in these towns remove snakes and other reptiles when requested by members of the public. Reptile removal records for Kununurra were consistently kept from 2006 prior to cane toad invasion in 2010, through to 2017, encompassing both pre- and post-invasion periods (Fig. 2b). Wildlife officers are compelled to attend snake callouts for reasons of public safety, so data can be considered representative of snake occurrences in the town. Records included date and time of removal, the officer’s name who attended, the location/address, the species and size (length) of the animal removed. Data are presented as monthly counts for analysis. In Katherine, snake removal records were available from 1998 to 2008 and covered the pre-invasion (1998-2000) to post-cane toad invasion (2001-2008) periods. Darwin snake removal data were available more sporadically during nine non-consecutive years encompassing pre- and post-invasion by toads. Annual species counts were made for 1997, 1998 and 1999 prior to toad invasion. Records including attending officer, date, time, address, actions and species were kept for the post-toad period in 2005, and then annually from 2011 to 2015. Despite the potential for species and habitat bias in the wildlife removed from urban/rural environments in towns [44], we assume that the reptile data represent surrounding savannas because of the small size and isolation of these towns within the vast expanse of uninhabited savannas across the whole of northern Australia (Fig. 2a). This assumption is supported by records of species identities, as most snakes and other reptiles removed were common species characteristic of northern Australian savanna assemblages [45].

### Fauna surveys

To obtain additional information on predators, and to sample populations of potential prey, we conducted fauna surveys at 15 sites in Mirima NP, Kununurra, from 2006 to 2017 (see Fig. 2a). This period encompassed both pre- and post-cane toad invasion. Mammal trapping data from Elliott and pitfall trap surveys were available for 23 months, though not all sites could be surveyed every month due to logistical constraints (Table S1). Surveys at sites m, n and o in Jul 2006, Jan and Sep 2007, May 2008, Mar and Apr 2017 (Table S1) used 50 × 50 m quadrats and 20 Elliott traps (alternating large 15 × 15.5 × 46 cm and medium 9 × 10 × 33 cm traps) around the perimeter and 10 pitfall traps (20 cm diameter, 60 cm deep) placed along two parallel drift fence-lines [34, 26]. Mammal surveys at sites a, b, c, d, e, f, g, h, i, j, k and l from Mar 2010 to Apr 2017 (Table S1) used a 40 × 100 m grid with 18 Elliott traps (alternate 9 large and 9 medium traps) placed 20 m apart in the grid and 4 pitfall traps (29 cm diameter 40 cm deep) placed at each corner with 4 shallow trenches (5 to 15 cm deep) directing animals into traps [46]. Mammal surveys occurred for either 4 or 7 nights. All mammals were identified to species, weighed, head and body length measured, and marked prior to release (permanent marker on ear). Recaptures were not counted. Mammal data are presented as total mammals per 1000 trap nights to standardise them; low numbers for individual mammal species represented in the surveys (Table S2) precluded species analyses.

Funnel trapping was used to survey reptile, frog and invertebrate assemblages during 25 months between Jul 2008 and Apr 2017 (Table S1). A 40 × 100 m survey grid was used with 18 funnel traps (18 cm × 60 cm) placed 20 m apart within the grid (Radford & Fairman, 2015). Funnel traps were placed in the middle of a 6 m long shallow trench (5 to 15 cm deep) to attract and direct animals into traps. All reptiles and frogs were identified to species and snout vent and tail length measured; animals were marked (permanent marker pen) prior to release to establish recaptures. Insects (> 5 mm long) were identified to Order or Family, and other invertebrates to Class or Order. Vertebrate species and invertebrate taxa were categorized according to trophic roles for analysis depending on their diets [47-52, 45]. Counts of reptile, frog and invertebrate species/taxa per trap session were recorded and used in analyses. For mammal, reptile, frog and invertebrate taxa we consider all Mirima survey sites, irrespective of survey methodology, to be sampling replicates for the sake of the analyses due to similarities in productivity and geology (sand or sandstone), vegetation (hummock savanna woodland/shrubland) and fauna assemblages.

### Arrival of cane toads

The arrival month of cane toads in Kununurra was set as the date when animals were first placed in bins at drop-off points by members of the public. The first records of toads in Kununurra were in April 2010. A second arrival date was set at the Mirima NP fauna survey sites adjacent to Kununurra when the first toads appeared in survey traps in April 2011 [46]. Cane toads first arrived in Katherine in 2001 and in Darwin in 2005 (T. Parkin, G. Gillespie, unpublished data).

### Statistical modelling

Species with fewer than 20 records were not included in modelling analyses. We used tscount [53] in R version 3.5.1 [54] to fit generalised linear models to our time series count data for each species or species group at different sites, i.e. integer-valued GARCH log-linear models with logarithmic link. In this way, the conditional mean could be linked to potential covariates (e.g. rainfall, temperature, etc., Table S2) and past values or past observations (i.e. previous means). We captured short range serial dependence using a first order autoregressive term on the previous observation (beta_1) and yearly seasonality using a 12^th^ order autoregressive term (alpha_12). Either a Poisson, or in the case of over-dispersion, a negative binomial conditional distribution, was chosen. Model fit and assessment were based on probability integral transform histograms, the autocorrelation function (ACF) of response residuals, and a cumulative periodogram of Pearson residuals. Using backward stepwise elimination, covariates were excluded on improvement in the model Akaike Information Criterion and only significant covariates (of those listed in Table S2) were included in final models (Table 1). Autoregressive terms were adjusted if the ACF plot indicated subsequent autocorrelation beyond beta_1 and alpha_12.

**Table 1.**
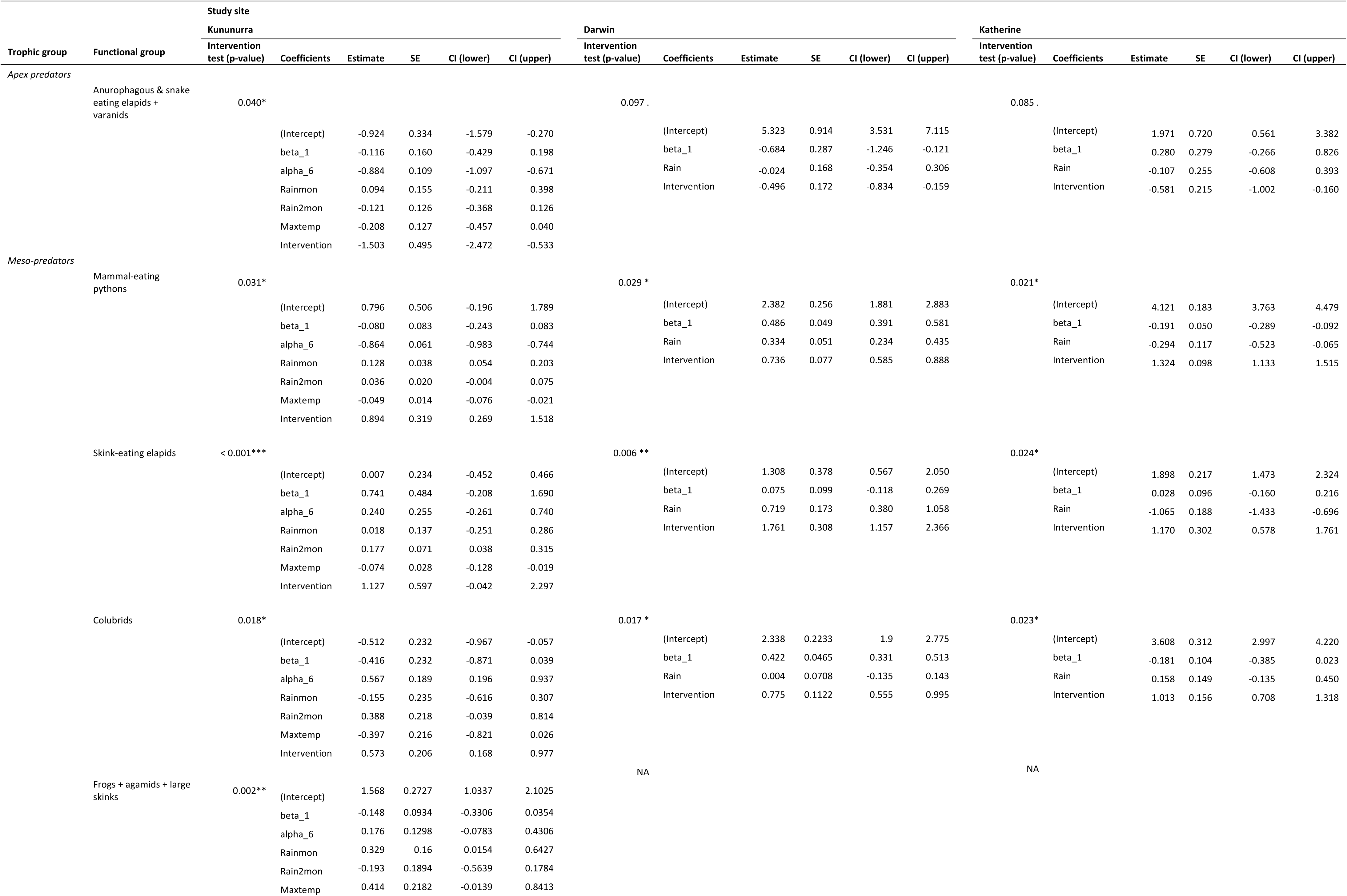
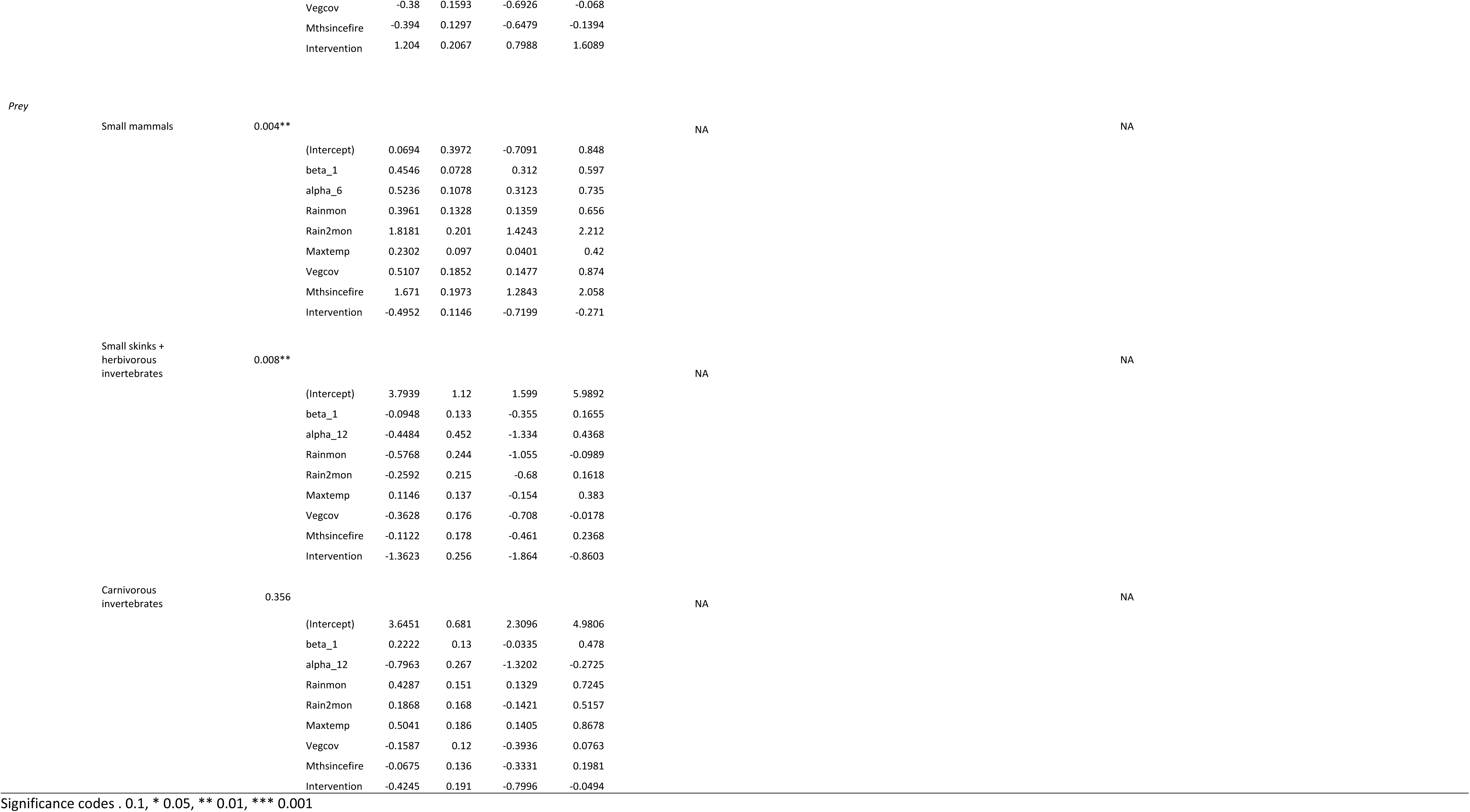
Level of significance (*p*-values) of intervention tests conducted at time of cane toad (*Rhinella marina*) invasion (Kununurra 2010, Darwin 2005, Katherine 2001) after fitting time series generalised linear models (INGARCH log-linear models with logarithmic link) on occurrence records of functional groups of apex predators, meso-predators and prey trophic levels at different sites.

To identify shifts in faunal count data post-cane toad invasion, we used intervention analyses (R package tscount) [53], where intervention, as defined by Fokianos & Fried [55], was included as a covariate in each model. This covariate included an integer vector giving the time when the intervention effect occurred (tau); in our case, tau differed for each sample site depending on when toads arrived or the detection of a lag in intervention effect. The intervention covariate also included a numeric vector with constants specifying the type of intervention (delta (δ), for 0 < δ < 1 the effect decays exponentially and for δ = 1 there is a persistent effect of the intervention after its occurrence). We chose δ = 1 as there was a persistent intervention effect after toad invasion and we were testing for a permanent shift post-intervention. Significance of the intervention effect was assessed for each species using the corresponding confidence intervals of intervention covariate coefficient estimates, and for trophic groups using an intervention test (valid only for long time series or large sample sizes) to test for intervention of type δ = 1 at the time of cane toad invasion.

## RESULTS

### Reptile removal and fauna survey data

Apex predatory species, including five species of elapid snakes (n = 364) and six species of varanids (n = 42); meso-predatory species/taxa, including 23 snake (n = 6584), nine lizard (n = 561) and seven frog species (n = 487); and prey species/taxa, including five mammals (n = 104), six lizards (n = 599) and six invertebrate taxa (n = 1221) were recorded during removals and surveys (Table S2). During Kununurra removals, 328 snakes and reptiles were recorded during 130 consecutive months from Mar 2006 to Dec 2017 (Table S1). In Katherine, 1430 snakes were recorded during 11 years of callouts (1998 to 2008), and in Darwin 5168 snakes and lizards were recorded during nine non-consecutive years (1997-1999, 2005, 2011-2015) (Table S1, S2). In Mirima NP, 2932 reptiles, frogs and invertebrates were recorded during 25 non-consecutive monthly survey periods from 2008 to 2017 (Tables S1, S2). Small mammals were recorded during 23 survey months in Mirima NP between 2006 and 2017 (Table S1, S2).

### Responses to toad invasion among predator and prey groups

As predicted under the conceptual model (Fig. 1), apex predators declined significantly after cane toad invasion, almost all meso-predators increased, and most prey groups – including small mammals – also declined based on intervention tests (Table 1, Figs 3, 4). For combined apex predators in Kununurra, the largest GLM coefficient estimates were for intervention, indicating a strong impact of toad invasion relative to other explanatory variables (Table 1). The strongest additional explanatory variable for apex predators was a 6 month temporal auto-correlative effect indicating a seasonal influence on predator numbers (Table 1, Fig. 3a, Fig. 4a). Intervention (toad invasion) was also significant (marginally) among apex predators in both Darwin and Katherine, with rainfall and monthly autocorrelative terms supported as additional explanatory variables in the model (Table 1).

**Fig. 3.**
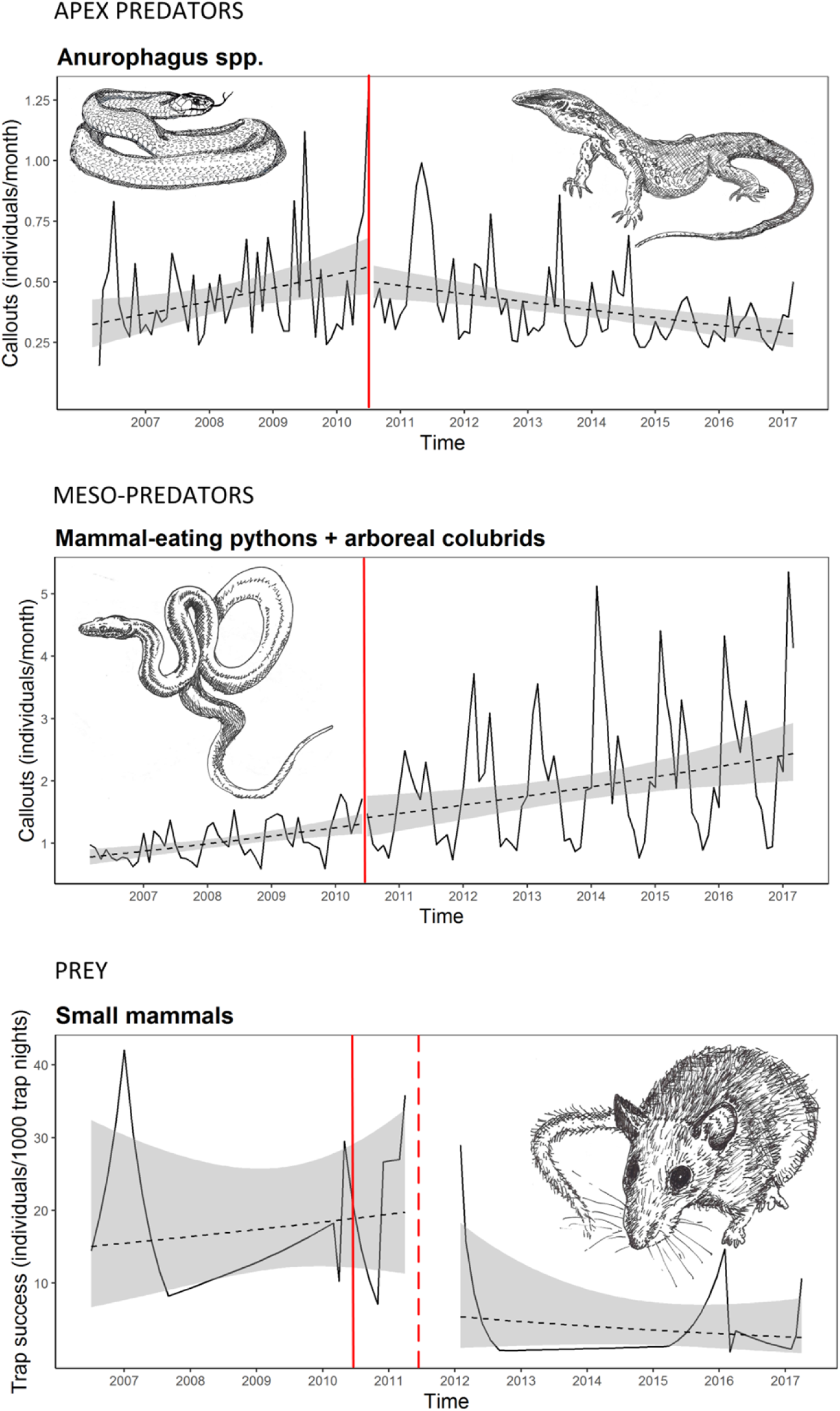
Modelled trends in occurrence before and after cane toad (*Rhinella marina*) invasion for a) apex predators (large anurophagous reptiles and elapids) from Kununurra snake callout records, b) meso-predators (mammal-eating pythons and arboreal colubrids) from Kununurra snake callout records, and c) the small mammal prey group from Mirima fauna surveys. The vertical solid red line indicates the arrival date of toads in Kununurra in 2010 and the dashed red line indicates when toads arrived at fauna survey sites in 2011.

**Fig. 4.**
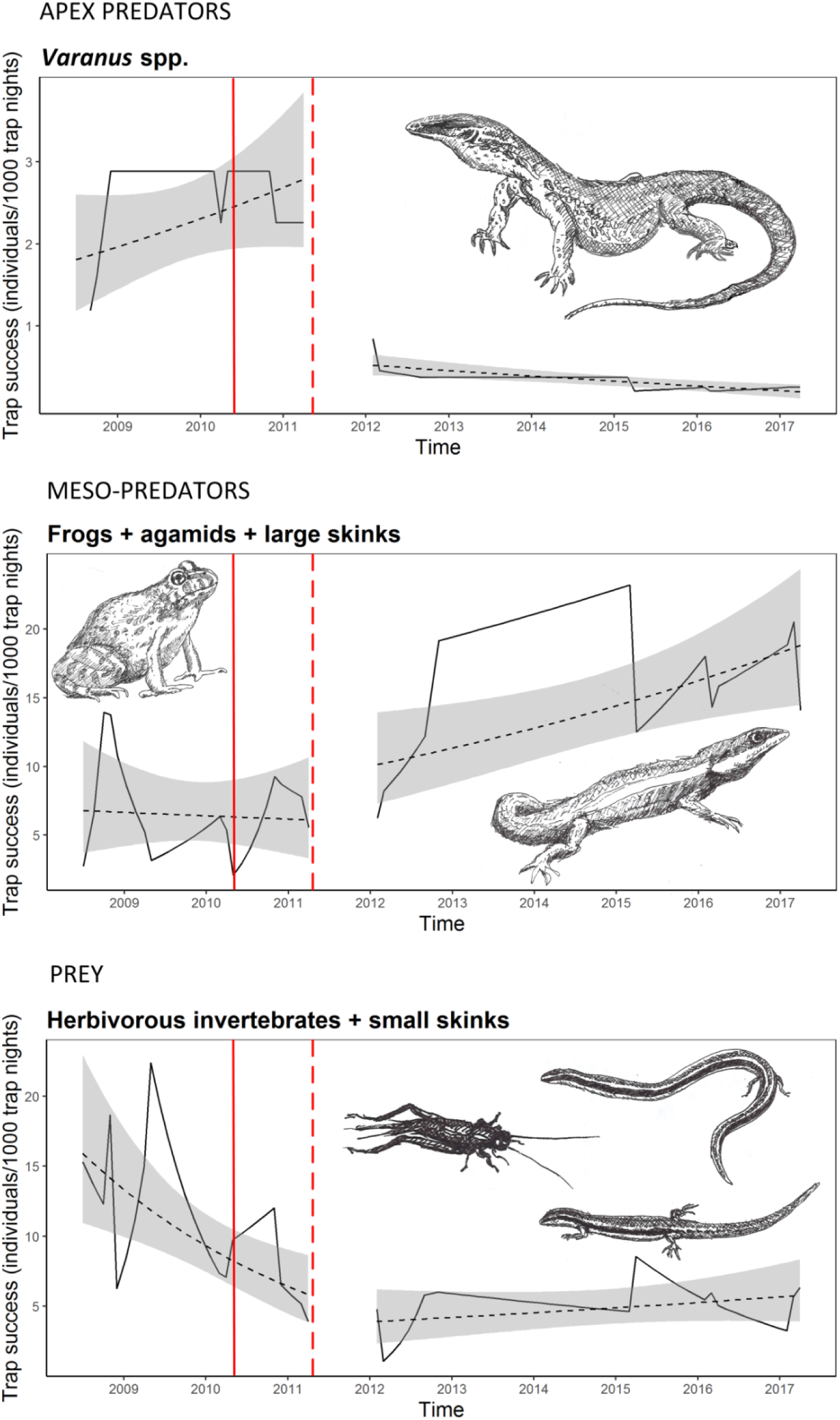
Modelled trends in occurrence (trap success) before and after cane toad (*Rhinella marina*) invasion for apex predators (large varanid reptiles, *Varanus* spp.), meso-predators (frogs, agamids and large skinks), and prey groups (herbivorous invertebrates and small skinks) from Mirima fauna surveys. The vertical solid red line indicates the arrival date of toads in Kununurra in 2010 and the dashed red line indicates when toads arrived at fauna survey sites in 2011.

Four of the five identified meso-predator groups, including the mammal-eating pythons (Fig. 3b), skink-eating elapids, colubrid snakes and combined frogs, agamids and large skinks (Fig. 4b), showed significant increasing intervention responses (Table 1). Intervention responses had higher coefficient estimates than for all other explanatory variables for meso-predators, indicating that toad invasion was the strongest predictor of change among these groups (Table 1). All three meso-predatory groups from Katherine and Darwin (mammal-eating pythons, skink-eating elapids and colubrid snakes) also had significant intervention responses (Table 1). The smallest of the identified meso-predators, *Eremiascincus*/*Heteronotia* (Table S2) had no significant intervention response (Table 1). Additional strongly supported explanatory variables for meso-predatory snakes (pythons, small elapids and colubrids) included seasonal auto-correlation (alpha_6) and rainfall, and for the frog/agamid/large skink group included rainfall in the previous month, maximum temperature, vegetation cover and time since fire (Table 1).

Among prey groups, small mammals showed a significant negative response to intervention (invasion) (Table 1, Fig. 3c). Other prey groups (e.g. herbivorous invertebrates (< 3 cm), small skinks, Table S2) also declined significantly post-invasion (Table 1, Fig. 4c). Exceptional among savanna prey groups were larger carnivorous invertebrates (3-12 cm, Table S2) which did not show a significant intervention response (Table 1). In addition to intervention, the strongest explanatory variables for small mammals were rainfall in the previous 2 months, and months since fire (Table 1). Herbivorous invertebrates and small skinks responded most strongly to rainfall in the previous month, seasonal auto-correlation and vegetation cover (Table 1). Large carnivorous invertebrates responded most strongly to seasonal auto-correlation, maximum temperature and rainfall in the previous month (Table 1).

Species-specific apex, meso-predator and prey responses generally conformed to responses predicted in the conceptual model (Fig. 1) with a few minor exceptions (Table S4). Among meso-predatory species, the skink *Eremiascincus isolepis* and gecko *Heterontia binoei,* in Kununurra, showed no intervention responses. These were among the smallest of the meso-predators (13 cm and 9 cm, Table S2) The lesser black whip snake, *Demansia vestigiata*, in Katherine, uniquely among small skink-eating elapids, showed a significant negative intervention response (Table S4). Among small skinks, *Carlia* spp. showed a positive intervention response which was the opposite to small skink responses overall (Table S4). *Carlia* spp. were the largest among the small skinks (ca. 11 cm) with others in the group < 10 cm long (Table S2).

## DISCUSSION

Small mammal declines in northern Australia have not previously been temporally or spatially linked with the arrival of cane toads [34, 20-21], nor to cascading impacts among reptilian predators [14, 19]. To some extent this may have arisen due to small numbers of the larger reptilian predators usually trapped during standard fauna surveys leading to difficulties in detecting trends coinciding with changes among smaller more numerous species including mammals. In this study we present simultaneous data on large reptilian predator assemblages using novel wildlife removal data alongside standard fauna monitoring data collected at the same time and place. These combined data reveal multiple and pervasive ecosystem-wide trends coincident with cane toad invasion which have not previously been apparent from standard surveys alone. The trends identified in this study are consistent with the hypothesis that cane toad invasion initiates ecosystem-wide trophic cascades (Fig. 1), as suggested by Doody et al. [14] and Feit et al. [19]. These trophic cascades include the functional loss of keystone apex reptilian predators including large-gaped elapid snakes [42] and large varanid lizards (*Varanus* spp); a meso-predatory release of smaller-gaped predominantly mammal- and skink-eating snakes (pythons, colubrids and small elapids) [15] and invertebrate-eating predators (e.g. frogs and agamid and large scincid lizards); and finally a decline among savanna prey groups including small mammals, smaller skinks and invertebrates, resulting from increased predation pressure by meso-predators (Fig. 1). Although previous studies have speculated that there may be a link between toad-driven changes to predator assemblages and small mammal declines [21], this is the first study to empirically link temporal and spatial data on toad invasion, reptilian predator assemblage change and small mammal declines.

Although the conceptual model presented here (Fig. 1) seems plausible, is partially supported by literature [39, 15, 14) and we have temporal and spatial links between mammal declines and toad invasion from Kununurra (Table 1), it is unclear if observed historical patterns of mammal decline align with sequential timing of toad invasions across northern Australia? Unfortunately monitoring programs for large reptilian predators and savanna fauna generally were not widespread, co-ordinated or sometimes even initiated prior to 2001-2005 [56] when cane toads first appeared in the Katherine/Darwin region (Fig. 2a). Therefore, there is little quantitative evidence to link mammal declines, their timing and/or their associated threatening processes [56]. However, it is clear that some northern mammal declines pre-date cane toad arrival. Mammal declines reported at mainland Northern Territory sites up to 2005 [23-24, 56-58], pre-2010 in the Kimberley region of Western Australia [22, 25-26], or up to the present on Melville Island [59] cannot be attributed to cane toad invasion because the invader had not yet arrived at these locations (Fig. 2a). However, many of these pre-toad declines affected only some mammal species or groups [22, 26, 56-57, 60-61], were subtle and relatively difficult to detect [23-24] or were based on few temporal data points [59], making it difficult to interpret changes as decline rather than as natural population variability.

In contrast to the above changes, more recent mammal declines post cane toad invasion in Kakadu and elsewhere in the Northern Territory since 2005 [34, 20, 56, 36, 21, 58] have been pervasive across the entire suite of critical weight range mammal species (mean adult body weight 35 - 5500 g) [62], have involved dramatic population collapses to levels almost beyond detectability, and have been relatively well documented. In addition, mammal abundance ahead of the cane toad invasion in the Kimberley has remained relatively high and stable throughout the same period (e.g. mean trap success 7.24%, as per Radford et al. 2014), until the declines noted in this study in the eastern edge of the Kimberley after cane toads arrived in 2010. In contrast, mammal abundance behind the cane toad front in the Northern Territory has remained consistently very low ever since invasion in 2005 (mean trap success < 1% trap success) [34, 58]. These data collectively support the notion that factors other than cane toads (e.g. cats and disturbance regimes) [4, 21] have been involved in driving small mammal declines across north western Australia pre cane toad invasion, but also that cane toad arrival has led to a recent increase in the pervasiveness of mammal assemblage-wide collapses on top of the previous declines.

Current thinking pertaining to small mammal declines in northern Australia centres on feral cat predation as a key driver [4, 20-21, 27-28, 31]. However, the cat and cane toad hypotheses are not incompatible and may act as complementary (and cumulative) drivers of mammal declines. The role of cats in northern Australia is seen as an extension of historical nationwide cat- and fox-driven mammal declines and extinctions, especially in the arid zone [4, 32]. However, the cat hypothesis relies on interactions with other factors to be a tenable explanation for northern mammal declines [20-21, 36]. This is because cats apparently coexisted with savanna mammals for over a century before recent north Australian declines [33]. Cat predation is known to interact with high intensity fire regimes to concentrate cat hunting activity [28, 30], and this increases mortality in local mammal populations [29, 31]. However, cat predation pressure is also thought to be influenced by apex predators, in particular the dingo (*Canis dingo*) [4-6]. High density of dingoes in high rainfall, high productivity areas of the Kimberley has been argued to reduce predation impacts by meso-predatory cats [5, 26]. However, cats are also known anecdotally to be depredated by large reptilian predators (and also to eat reptiles) [63]. As meso-predators, cats too may benefit from cane toad driven declines of apex reptilian predators, similar to those recorded here among meso-predatory snakes, lizards and frogs (Table 1). Future research is required to examine interactions between cane toad invasion, reptilian predator cascades and their interactions with cat impacts on small mammals if a greater understanding of mammal declines in northern Australia is to be achieved.

The conceptual model presented in this study has empirical support, highlighting the timing and apparent strength of observed trends associated with cane toad invasion. However, raising conceptual models/hypotheses to explain observed patterns is only one step in the process of establishing a model’s efficacy. The next step is for the conceptual model to be subject to tests, or falsified, to enable us to evaluate further if the hypothesis provides a tenable explanation for mammal declines relative to other hypotheses. Future research is needed on meso-predatory snake densities and predation rates to test whether predation by these reptilian predators is sufficient compared to that of cats to cumulatively drive mammal declines. Recently, estimates of cat densities and predation rates were made across the continent [63-66], including the Kimberley region [29]. Equivalent estimates are not available for reptilian predators and their impacts on mammals. What fragmentary data we have on snake densities and home ranges [67-70] suggest much greater densities of snakes than for cats and also much smaller home ranges. This means that even if snake ingestion rates are much lower than for cats, they may cause comparable overall predation pressure.

We know that snakes in some cases can have very large impacts on mammalian and avian assemblages. These include one meso-predatory snake from this study (e.g. *B. irregularis*) [16]. In addition, there is information from a cat exclosure experiment in northern Australia [35] that showed similar predation by pythons on savanna rodents to that by cats. However, the hypothesis that reptilian predation could be equivalent to that of feral cats, and the possibility that cumulative impacts could be substantial, needs to be tested more widely across Australian savanna landscapes if we are to establish its plausibility in playing part in regional mammal declines.

Another test to validate the role of cane toad cascades in driving mammal declines, is whether ongoing toad invasion across the Kimberley leads to rolling changes among reptilian (and mammalian) predator assemblages and to continuing mammal losses. Following initial cane toad arrivals in Kununurra/East Kimberley in 2009/2010, cane toads have now spread to Purnululu National Park in the south east Kimberley (ca. 2012), to Drysdale River National Park in the north Kimberley (ca. 2016), and to the far north Kimberley at Mitchell River National Park (ca. 2019) (Fig. 2a). If cane toad initiated cascades are a key factor driving mammal declines, we should expect mammal monitoring programs at these locations to reveal further declines within five years to one decade following toad arrival. Already the limited data from Purnululu National Park (Fig. 2a) indicates a ca. 90% decline in mammal trap success following toad invasion, with pre-toad trap success recorded at 1.4 % in 1989 [71], 1.5 % in 2004/2005 [72] and 3.7% in 2008 [26] prior to toads; at 2.5 % one year following cane toad arrival in 2013 (Fig. 2a); and then down to 0.42 % and 0.25 % in 2016 and 2017 four and five years post-invasion (I.J. Radford and B. Corey, unpublished data). Documentation of sequential reptilian predator changes associated with mammal declines and toad invasion in the wake of cane toad invasion fronts moving across the Kimberley would provide further empirical evidence for the role of cane toads in driving recent mammal declines. Ideally, these empirical studies of temporal and spatial changes coincident with cane toad invasion would be accompanied by experimental exclosure studies, similar to those conducted for cat impacts on mammals [27, 73, 58, 35], to test the plausibility and magnitude of reptilian predator impacts on mammals in both pre- and post-invasion savanna ecosystems.

This study joins others in highlighting the potential importance of reptilian predators, and reptilian meso-predator release, in the functioning of Australian and global ecosystems [1, 14, 16-17, 74]. One of the meso-predatory species implicated here in driving mammal declines, *Boiga irregularis*, is already documented as having driven catastrophic declines and extinctions of an entire avian forest assemblage, as well as small mammals, on the island of Guam [16]. It is perhaps not surprising that, in a continent with very high reptilian diversity [74-75] and several reptilian niche equivalents of mammalian predators elsewhere [18], as well as extensive pre-historical extinctions of most large mammal species [4], that reptile predators play such an ostensibly prominent role in Australian ecosystems.

## Data Accessibility Statement

Data from Western Australia and owned by the Department of Biodiversity, Conservation and Attractions can be made available upon request to the first author. Data owned by separate custodians including the Northern Territory’s Flora and Fauna Division, Department of Land Resource Management, or Rick Shine from the University of Sydney, would have to be requested separately direct to custodians.

## Supporting Information

**Table S1.** Mirima NP small mammal (Elliott traps, pit fall traps) and reptile, frog and invertebrate (funnel traps) fauna survey site numbers, trap effort and site names during months when surveys were conducted between 2006 and 2017. **^1^**Denotes the first month when cane toads (*Rhinella marina*) were recorded in drop off points in Kununurra (Intervention 1). **^2^**Denotes when toads first appeared in survey records at Mirima sites (Intervention 2).

| Year | Month | Mammals |  | Reptiles, Frogs, Invertebrates |  | Site names |
| --- | --- | --- | --- | --- | --- | --- |
|  |  | No. sites | Trap effort | No. sites | Trap effort |  |
| 2006 | Jul | 3 | 336 | - | - | m,n,o |
| 2007 | Jan | 2 | 224 | - | - | m,n |
| 2007 | Sep | 1 | 112 | - | - | o |
| 2008 | May | 1 | 112 | - | - | o |
| 2008 | Jul | - | - | 3 | 378 | a,b,c |
| 2008 | Oct | - | - | 2 | 252 | a,b |
| 2008 | Nov | - | - | 6 | 756 | c,d,e,f,g,h |
| 2008 | Dec | - | - | 2 | 252 | i,j |
| 2009 | Apr | - | - | 7 | 882 | a,b,d,e,f,g,j |
| 2009 | May | - | - | 3 | 378 | c,h,i |
| 2010 | Mar | 7 | 432 | 7 | 504 | a,b,d,e,f,g,h |
| 2010<sup>1</sup> | Apr | 2 | 288 | 2 | 144 | c,i |
| 2010 | May | 1 | 144 | 1 | 72 | j |
| 2010 | Nov | 8 | 704 | 8 | 576 | a,c,d,f,g,h,i,j |
| 2010 | Dec | 2 | 176 | 2 | 144 | b,e |
| 2011 | Mar | 7 | 616 | 7 | 504 | a,b,c,d,g,i,j |
| 2011<sup>2</sup> | Apr | 3 | 264 | 3 | 216 | e,f,h |
| 2012 | Feb | 6 | 528 | 6 | 432 | c,d,f,g,i,j |
| 2012 | Mar | 4 | 352 | 4 | 288 | a,b,e,h |
| 2012 | Sep | 6 | 528 | 6 | 432 | a,b,d,e,g,h |
| 2012 | Nov | 4 | 352 | 4 | 288 | c,f,i,j |
| 2015 | Mar | 6 | 528 | 6 | 432 | c,d,f,g,i,j |
| 2015 | Apr | 4 | 352 | 4 | 288 | a,b,e,h |
| 2016 | Feb | 6 | 528 | 6 | 432 | a,b,e,h,i,j |
| 2016 | Mar | 2 | 176 | 2 | 144 | d,g |
| 2016 | Apr | 4 | 352 | 4 | 288 | c,f,k,l |
| 2017 | Feb | 2 | 176 | 2 | 144 | c,d |
| 2017 | Mar | 8 | 644 | 8 | 616 | a,b,e,g,h,i,j,m |
| 2017 | Apr | 2 | 176 | 2 | 112 | n,f |

**Table S2.** Snake, reptile and mammal species represented in callout and survey records from Kununurra (KNX, Mirima), Katherine (KAT) and Darwin (DAR). Fauna species and taxa are grouped according to predatory hierarchy (apex- and meso-predators and prey) and functional groupings within the conceptual model (see Fig. X). Numbers in bold are those for which intervention analyses were undertaken. Dietary information for reptile and frog species/genera was taken from Cogger (2014) and from Menkhorst and Knight (2011) for small mammals. Size of each species is the mean length of animals captured during wildlife removals/surveys, or if not measured, maximum length as recorded in Cogger (2014). *Denotes the number of introduced cane toads collected at Kununurra drop-off points from April 2010 through to the end of 2013.

| Functional group | Species | Common name | SVL (cm) | Diet | KNX | KAT | DAR | Mirima (KNX) |
| --- | --- | --- | --- | --- | --- | --- | --- | --- |
| **APEX PREDATORS** |  |  |  |  |  |  |  |  |
| Large-bodied/gaped elapids | *Pseudechis australis* | King brown, mulga | 139 | reptiles including snakes, frogs, small mammals | **20** | **38** | **28** | 7 |
|  | *Pseudonaja nuchalis* | Western brown, gwardar | 106 | reptiles, small mammals, frogs | **28** | **68** | **144** | 7 |
|  | *Pseudonaja ingrami* | Ingram’s brown | 100 | vertebrates | - | 1 | - | - |
|  | *Pseudonaja textilis* | Eastern brown | 200 | small mammals, reptiles | - | 1 | - | - |
|  | *Pseudechis weigeli* | Pygmy, speckled mulga | 120 | vertebrates | - | **21** | - | 1 |
| Varanids | *Varanus gouldii* | Sand monitor, Gould’s monitor | 113 | reptiles, insects, birds, mammals, frogs, carrion | 6 | - | - | 16 |
|  | *Varanus acanthurus* | Spiny tailed monitor | 70 | lizards, insects | - | - | - | 6 |
|  | *Varanus mertensii* | Merten’s water monitor | 65 | fish, frogs, carrion | 4 | - | - | - |
|  | *Varanus mitchellii* | Mitchell’s water monitor | 35 | insects, fish, frogs | - | - | - | 1 |
|  | *Varanus scalaris* | Spotted tree monitor | 60 | insects, lizards | - | - | - | 1 |
|  | *Varanus tristis* | Black headed monitor | 60 | arboreal reptiles, insects | - | - | - | 2 |
|  | **All varinids** | goannas, monitors | 81 | reptiles, insects, birds, mammals, frogs, carrion | **10** | - | 6 | **26** |
| **MESO-PREDATORS** |  |  |  |  |  |  |  |  |
| Pythonids | *Antaresia childreni* | Childrens python | 60 | small mammals, birds, reptiles | **48** | **295** | **720** | 2 |
|  | *Aspidites melanocephalus* | Black headed python | 128 | small mammals, ground birds, reptiles, venomous snakes | **12** | **49** | **27** | - |
|  | *Liasis fuscus* | Water python | 141 | small mammals, other vertebrates | **24** | **51** | **637** | - |
|  | *Liasis olivaceous* | Olive python | 160 | small mammals, other vertebrates | **71** | **266** | **237** | - |
|  | *Morelia spilota* | Carpet python | 162 | small mammals, terrestrial vertebrates | - | - | **993** | - |
| Colubrids | *Boiga irregularis* | Night tiger, brown tree snake | 98 | small mammal, birds, eggs, lizards | 5 | **183** | **297** | - |
|  | *Dendrelaphis punctulatus* | Common tree snake | 112 | semi-arboreal frogs, birds | **74** | **168** | **1094** | - |
|  | *Stegonotus cucullatus* | Slatey grey snake | 94 | lizards, small mammals | - | - | **668** | - |
|  | *Tropidonophis mairii* | keelback | 52 | frogs, semi-aquatic | 6 | **59** | **117** | 1 |
| Small-bodied/gaped Elapids | *Acanthophis rugosus* | Northern death adder | 70 | lizards, small mammals | - | - | 1 | 1 |
|  | *Brachyurophis roperi* | Northern shovel-nosed snake | 25 | reptile eggs | - | 1 | - | 11 |
|  | *Cryptophis pallidiceps* | Secretive snake, small eyed snake | 50 | lizards, frogs | - | **21** | **54** | - |
|  | *Demansia olivacea* | Olive whip snake | 73 | lizards | - | 6 | 9 | 1 |
|  | *Demansia papuensis* | Greater black whip snake | 97 | lizards | **21** | **100** | **71** | 2 |
|  | *Demansia simplex* | Grey whip snake | 60 | lizards | - | 1 | 3 | - |
|  | *Demansia vestigiata* | Lesser black whip snake | 160 | lizards | - | **63** | - | - |
|  | *Furina ornata* | Orange-naped, moonsnake | 36 | small skinks | 9 | **30** | - | - |
|  | *Suta punctata* | Spotted snake | 38 | lizards | - | 5 | - | 3 |
|  | *Vermicella* spp. | Bandy bandy | 50 | blind snakes | - | 2 | - |  |
| Acrochordids | *Acrochordus arafurae* | Arafura file snake | 150 | fish | - | 2 | 2 | - |
| Typhlopids | *Ramphotyphlops* spp. | Blind snakes | 30 | ants, termites | - | - | 1 | 2 |
| Homalopsids | *Fordonia leucobalia* | Crab eating snake | 100 | small fish, crabs | - | - | 3 | - |
| Agamids | *Chelosania brunnea* | Chameleon dragon | 22 | invertebrates | - | - | - | 3 |
|  | *Diporiphora magna* | Yellow-sided two-lined dragon | 25 | invertebrates | - | - | - | 92 |
|  | *Diporiphora pindan* | Pindan two-lined dragon | 17 | invertebrates | - | - | - | 61 |
|  | *Lophognathus gilberti* | Gilberts dragon | 38 | invertebrates | - | - | - | 14 |
|  | **All *Diporiphora*** |  |  |  |  |  |  | **153** |
| **Large Scincids** | *Ctenotus inornatus* | Bar-shouldered ctenotus | 19 | invertebrates | - | - | - | 123 |
|  | *Ctenotus pantherinus* | Leopard ctenotus | 27 | invertebrates | - | - | - | 1 |
|  | *Ctenotus piankai* | Pianka’s ctenotus | 19 | invertebrates | - | - | - | 5 |
|  | *Ctenotus robustus* | Robust ctenotus | 24 | invertebrates | - | - | - | 248 |
|  | *Ctenotus tantillus* | Kimberley wedge snout ctenotus | 18 | invertebrates | - | - | - | 2 |
|  | **All *Ctenotus*** |  |  |  |  |  |  | **379** |
|  | *Eremiascincus isolepis* | Northern bar-lipped skink | 13 | invertebrates |  |  |  | 94 |
| **Gekkonids** | *Heteronotia binoei* | Binoe’s gecko | 9 | invertebrates |  |  |  | 107 |
| **Anurans** | *Crinia biligua* | Bilingual froglet | 4 | invertebrates | - | - | - | 18 |
|  | *Cyclorana* spp. | Large terrestrial frogs | 5 | invertebrates | - | - | - | 40 |
|  | *Litoria* spp. | Tree frogs | 4 | invertebrates | - | - | - | 22 |
|  | *Lymnodynastes convexiusculus* | Marbled frog | 4 | invertebrates | - | - | - | 13 |
|  | *Notadon melanoscapus* | Northern spadefoot frog | 4 | invertebrates | - | - | - | 9 |
|  | *Platyplectrum ornatum* | Ornate frog | 3 | invertebrates | - | - | - | **266** |
|  | *Uperoleia* spp. | toadlets | 2 | invertebrates | - | - | - | **131** |
|  | **All frogs** |  |  |  |  |  |  | **487** |
|  | *Rhinella marina* | Cane toad | 5 | invertebrates | 22623* | - | - | 205 |
| **PREY** |  |  |  |  |  |  |  |  |
| **Rodents** | *Pseudomys delicatulus* | Delicate mouse | 14 | seeds, plant matter, insects | - | - | - | 59 |
|  | *Pseudomys nanus* | Western chestnut mouse | 26 | grass stems | - | - | - | 10 |
|  | *Rattus tunneyi* | Pale field rat | 34 | vegetation | - | - | - | 22 |
|  | *Zyzomys argurus* | Common rock rat | 24 | seeds, plant matter, insects | - | - | - | 8 |
| **Small Dasyurids** | *Planigale ingrami* | Long tailed planigale | 12 | invertebrates, small vertebrates | - | - | - | 5 |
|  | **All small mammals** |  |  |  | - | - | - | **104** |
| **Small Scincids** | *Carlia* spp. | Rainbow skinks | 11 | small invertebrates | - | - | - | **234** |
|  | *Lerista griffini* | Stout sandslider | 9 | ants, termites | - | - | - | **269** |
|  | *Menetia greyii* | Dwarf skink | 6 | small invertebrates | - | - | - | 6 |
|  | *Morethia ruficauda* | Fire-tail skink | 8 | small invertebrates | - | - | - | 57 |
|  | *Notoscincus ornatus* | Ornate skink | 7 | small invertebrates | - | - | - | 31 |
|  | *Proablepharus tenuis* | Northern soil crevice skink | 6 | small invertebrates | - | - | - | 2 |
|  | **Tiny skinks (< 8 cm)** |  |  |  |  |  |  | **96** |
| **Large invertebrates** | Carnivorous invertebrates |  |  |  | - | - | - | **806** |
|  | Herbivorous invertebrates |  |  |  | - | - | - | **415** |
|  | Carabidae | Carabid beetles | 3 | invertebrates, small vertebrates | - | - | - | **379** |
|  | Chilopoda | Centipedes | 12 | invertebrates, small vertebrates | - | - | - | **318** |
|  | Blattodea | Cockroaches | 2 | detritus, plant material | - | - | - | **254** |
|  | Gryllidae | Crickets | 2 | plant material | - | - | - | **68** |
|  | Scorpiones | Scorpions | 5 | invertebrates | - | - | - | **44** |
|  | Araneae | Spiders | 2 | invertebrates | - | - | - | **61** |
|  | **All invertebrates** |  |  |  |  |  |  | **1221** |

**Table S3.** Covariates used in modelling of time series generalised linear models (INGARCH log-linear models with logarithmic link). Italics indicate name used in reporting of modelling results (Table 1, Table S3). All continuous variables were centred and standardised by dividing by two times the standard deviation [76].

| Parameter | Coding | Comment |
| --- | --- | --- |
| *beta_1* | As defined in R package tscount [53] | 1<sup>st</sup> order autoregressive term on previous observation in time series |
| *alpha_6, alpha_12* | As defined in R package tscount [53] | 6<sup>th</sup> or12<sup>th</sup> order autoregressive term on 6-monthly or 12-monthly (respectively) previous mean in time series |
| Intervention | As defined in R package tscount [53] | Intervention covariate for month of cane toad invasion in (1) Kununurra, May 2010; (2) Darwin, 2005; (3) Mirima, April 2011; (4) Katherine, 2001; (5) Katherine, 2005 |
| Maximum temperature<br>(*maxtemp*) | Continuous, centred, standardised | Mean maximum temperature (°C), monthly for Kununurra or Mirima, yearly for Darwin or Katherine |
| Monthly rainfall<br>(*rainmon*) | Continuous, centred, standardised | Mean monthly rainfall (mm) for Kununurra or Mirima |
| Two-monthly rainfall<br>(*rain2mon*) | Continuous, centred, standardised | Mean rainfall over previous two months (mm) for Kununurra or Mirima |
| Annual rainfall (*rain*) | Continuous, centred, standardised | Mean annual rainfall (mm) for Darwin or Katherine |
| Time since fire<br>(*mthsincefire*) | Integer, standardised | Number of months since last fire, included as covariate in Mirima models |
| Vegetation cover<br>(*vegcov*) | Integer, standardised | Percentage cover of sub-canopy shrub and herbaceous vegetation, included as covariate in Mirima models |

**Table S4.**
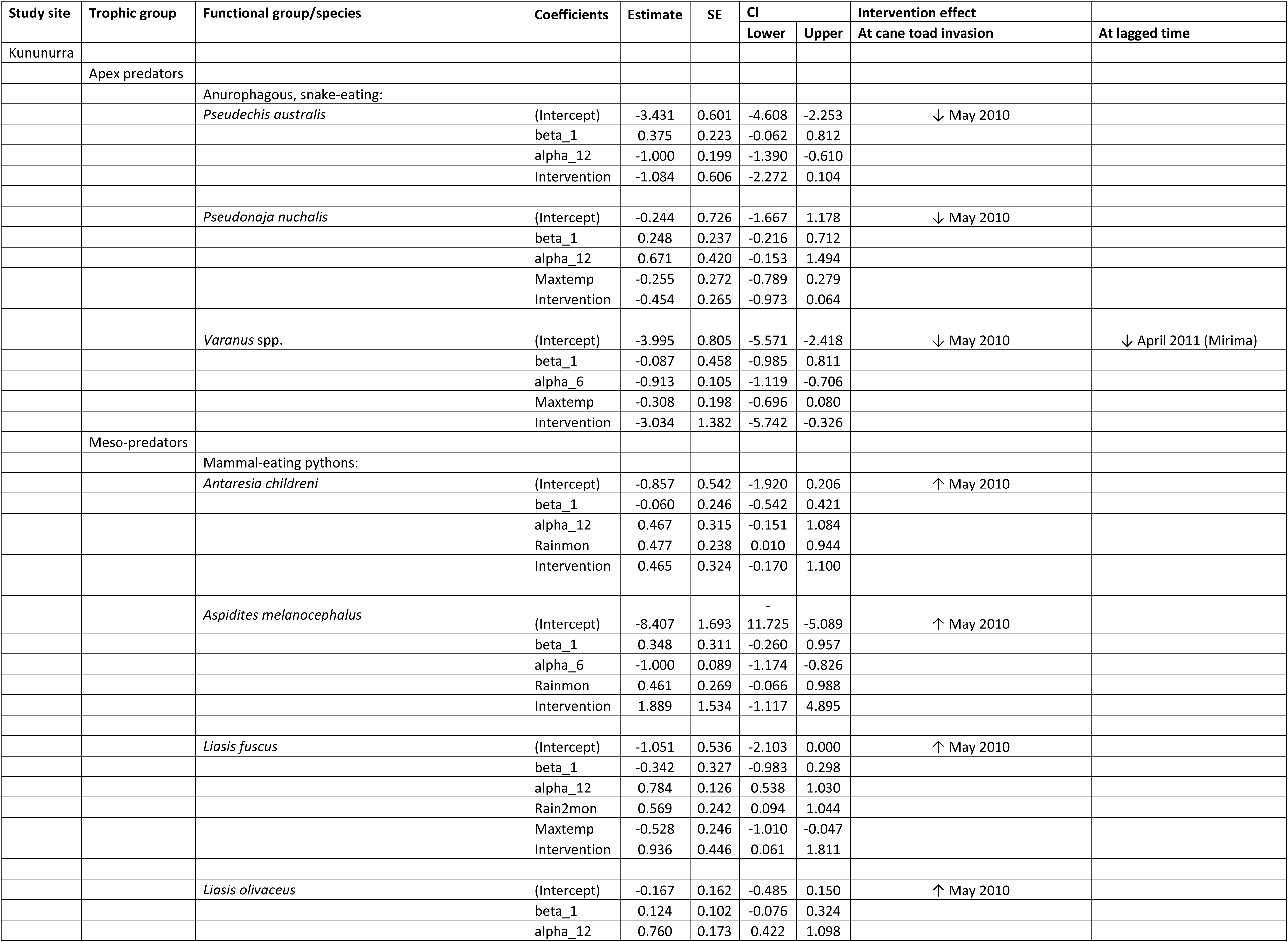
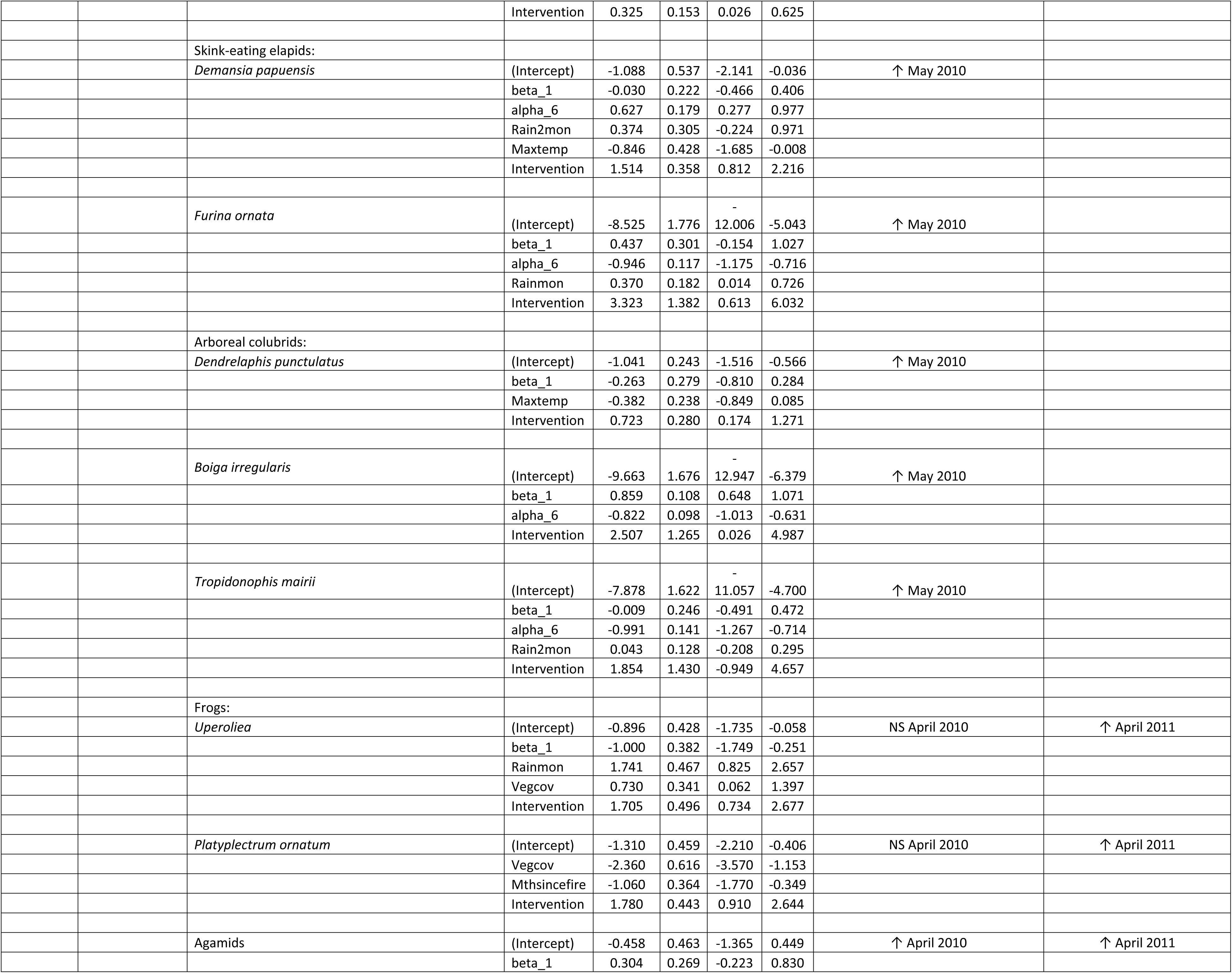
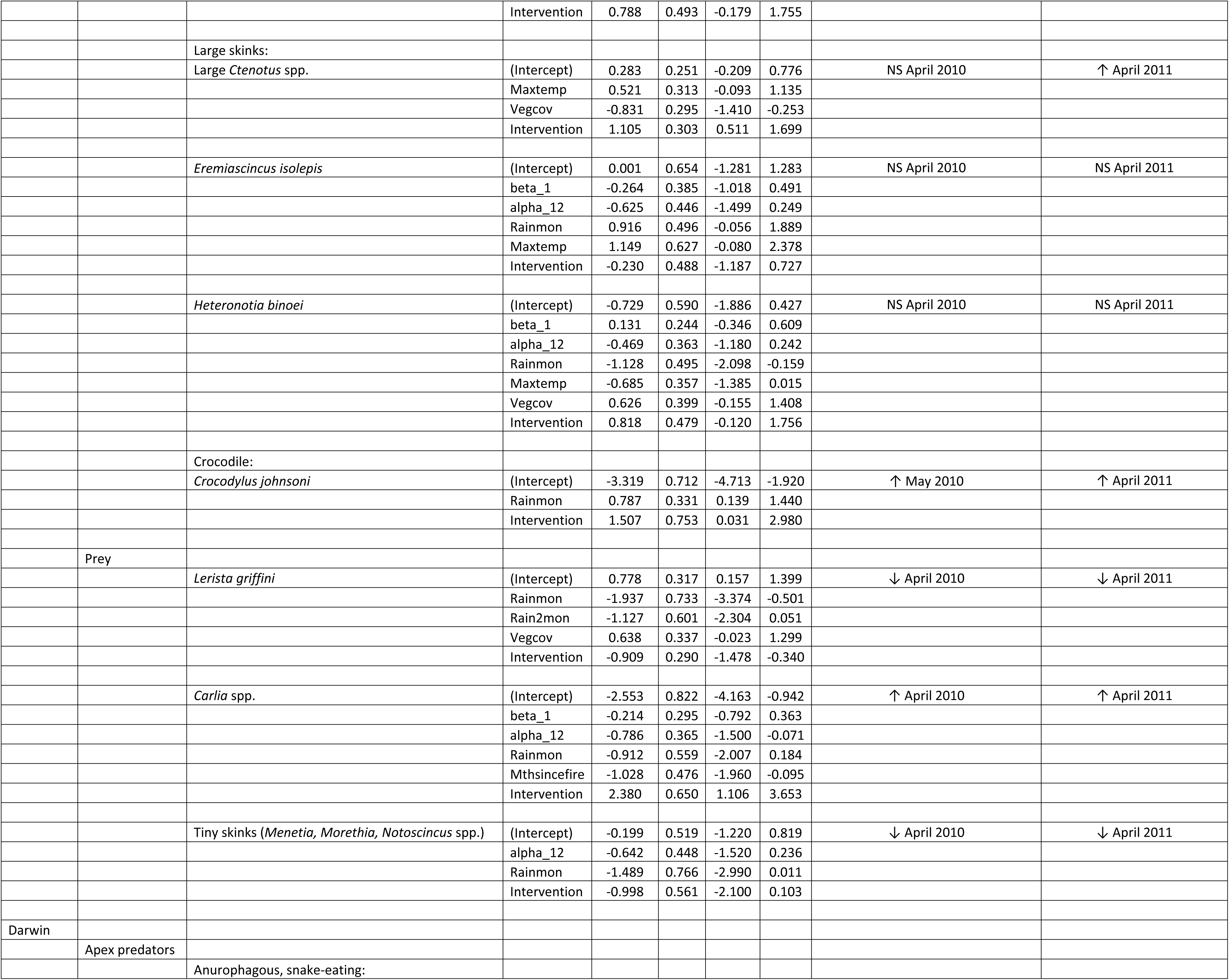
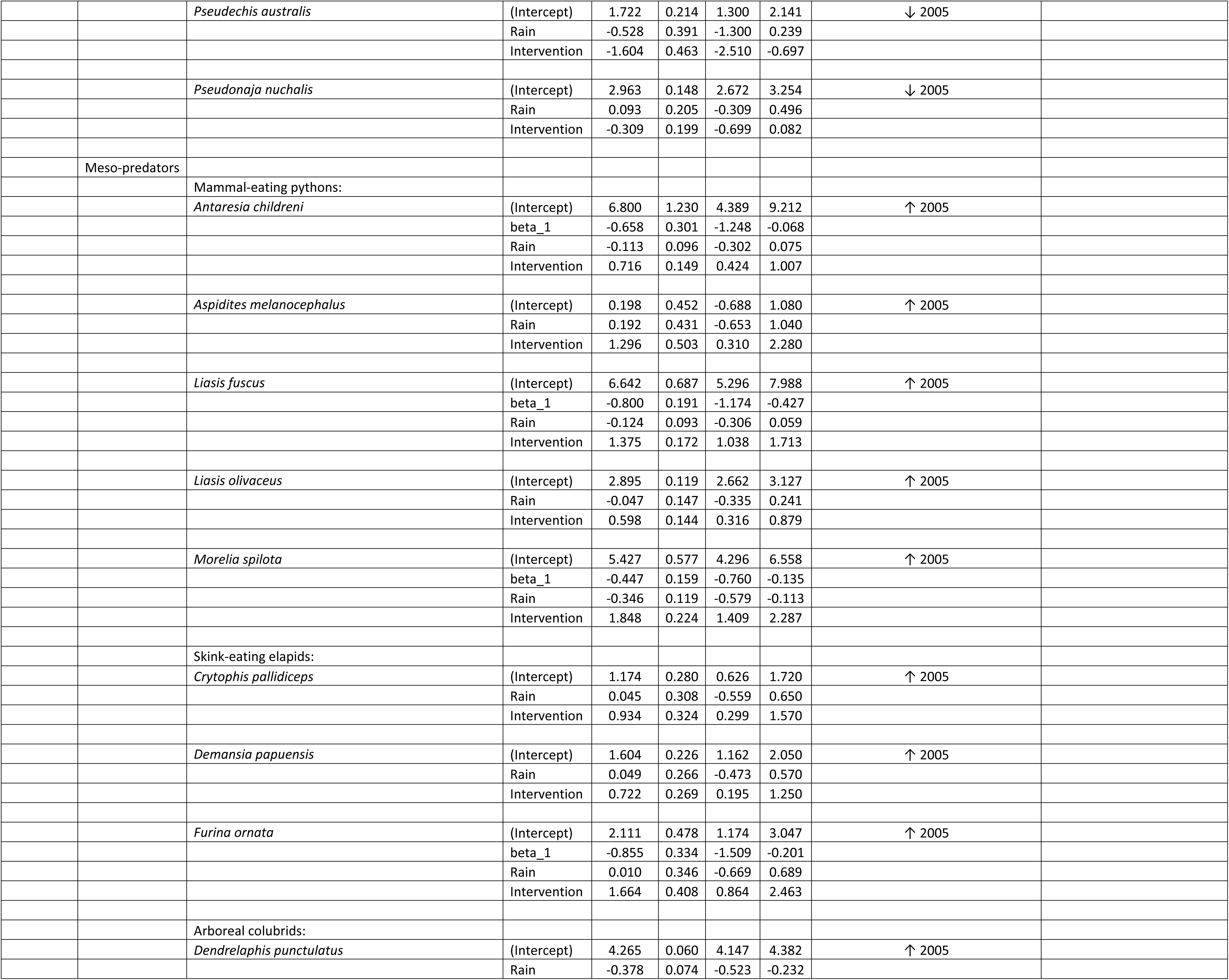
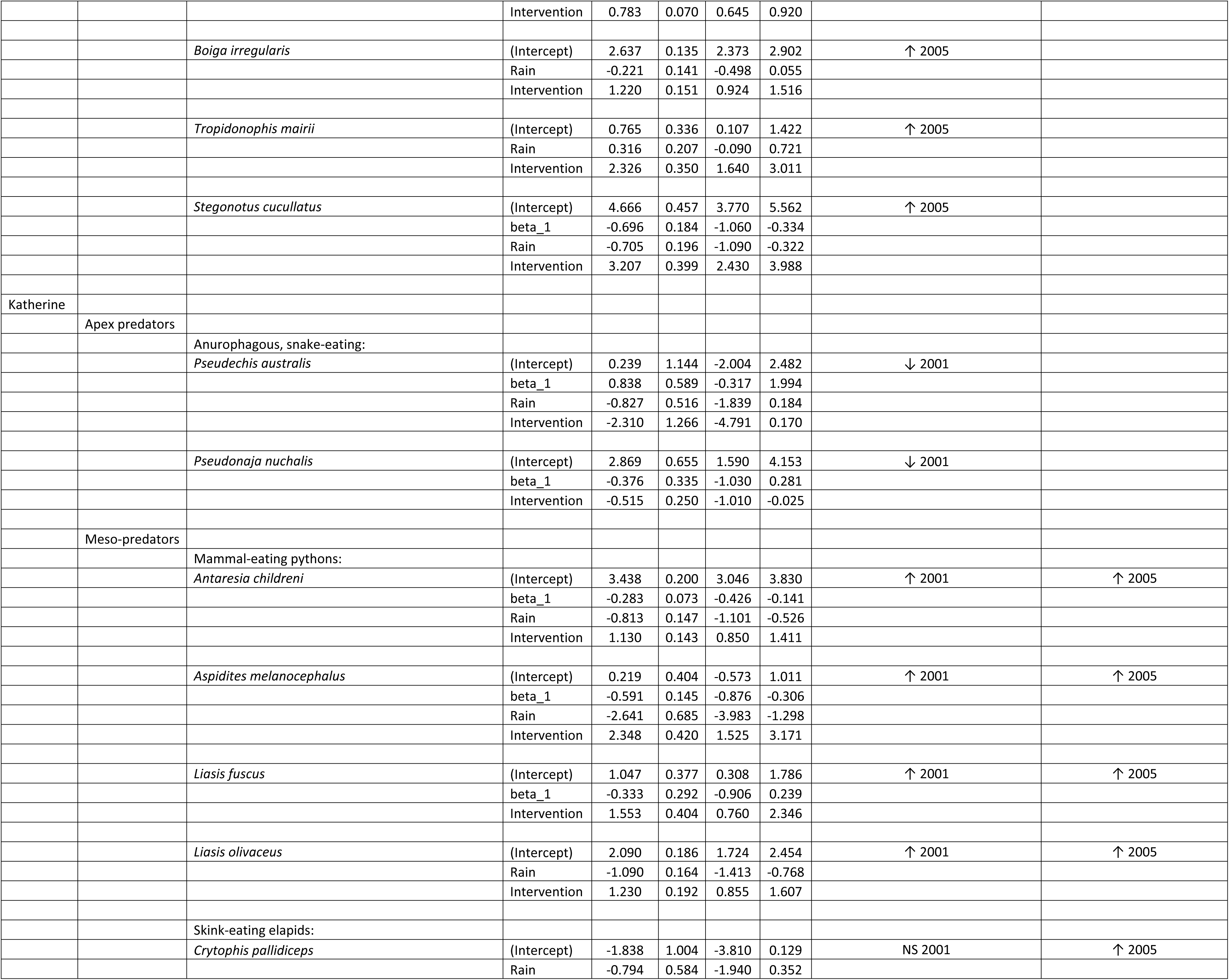
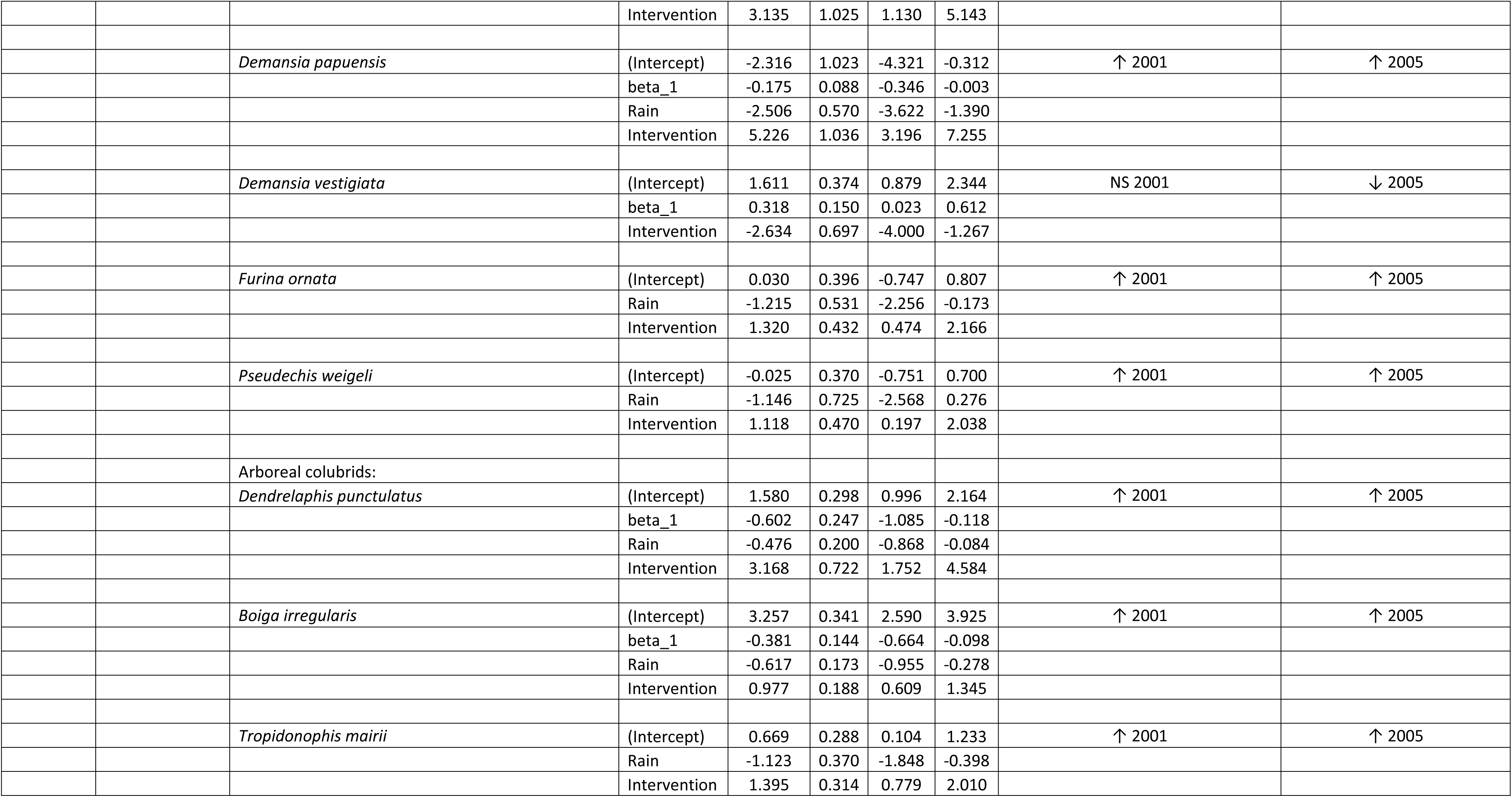
Time series generalised linear models (INGARCH log-linear models with logarithmic link) from occurrence records of species/functional/trophic groups of apex, meso-predators and prey trophic levels at different sites. Model coefficient estimates, standard error (SE) and 95% confidence intervals (CI) are shown. Intervention effect at the time of cane toad invasion (and/or at a lagged time after cane toad invasion) is indicated as significantly negative (↓), significantly positive (↑), or non-significant (NS) and the time of intervention effect is given. For definitions of variables see Table S2b.

## REFERENCES

1. Ritchie EG, Johnson CN. Predator interactions, mesopredator release and biodiversity conservation. Ecology Letters. 2009;12:982–998.

2. Garrott RA, Gude JA, Bergman EJ, Gower C, White PJ, Hamlin KL. Generalizing wolf effects across the Greater Yellowstone Area: a cautionary note. Wildlife Society Bulletin. 2005;33:1245–1255.

3. Allen BL, Engeman RM, Allen LR. Wild dogma: an examination of recent “evidence” for dingo regulation of invasive mesopredator release in Australia. Current Zoology. 2011;57:568–583.

4. Johnson C. Australia’s mammal extinctions: a 50,000-year history. Cambridge University Press. 2006.

5. Kennedy M, Phillips BL, Legge S, Murphy SA, Faulkner RA. Do dingoes suppress the activity of feral cats in northern Australia? Austral Ecology. 2012;37:134–139.

6. Letnic M, Ritchie EG, Dickman CR. Top predators as biodiversity regulators: the dingo *Canis lupus dingo* as a case study. Biological Reviews, 2012;87;390–413.

7. Newsome TM, Ballard G-A, Crowther MS, Dellinger JA, Fleming PJS, Glen AS, et al. Resolving the value of the dingo in ecological restoration. Restoration Ecology. 2015;23:201–208.

8. Leo V, Reading RP, Gordon C, Letnic M. Apex predator suppression is linked to restructuring of ecosystems via multiple ecological pathways. Oikos. 2018;

9. Levi T, Wilmers CC. Wolves–coyotes–foxes: a cascade among carnivores. Ecology. 2012;93:921–929.

10. Krebs CJ. The Ecological World View. University of California Press. 2008

11. Doody JS, Green B, Sims R, Rhind D, West P, Steer D. Indirect impacts of invasive cane toads (*Bufo marinus*) on nest predation in pig-nosed turtles (*Carettochelys insculpta*). Wildlife Research. 2006;33:349–354.

12. Doody JS, Green B, Rhind D, Castellano CM, Sims R, Robinson T. Population-level declines in Australian predators caused by an invasive species. Animal Conservation. 2009;2:46–53.

13. Doody JS, Castellano CM, Rhind D, Green B. Indirect facilitation of a native mesopredator by an invasive species: are cane toads re-shaping tropical riparian communities? Biological Invasions. 2013;15:559–568.

14. Doody SJ, Rhind D, Green B, Castellano C, McHenry C, Clulow S. Chronic effects of an invasive species on an animal community. Ecology. 2017;10:1002/ecy.1889.

15. Feit B, Letnic M. Species level traits determine positive and negative population impacts of invasive cane toads on native squamates. Biodiversity and Conservation. 2015;24:1017–1029.

16. Savidge JA. The Role of Disease and Predation in The Decline of Guam’s Avifauna (Extinction, Boiga Irregularis) Doctoral dissertation, University of Illinois at Urbana-Champaign. 1986.

17. Dorcas ME, Willson JD, Reed RN, Snow RW, Rochford MR, Miller MA, et al. Severe mammal declines coincide with proliferation of invasive Burmese pythons in Everglades National Park. Proceedings of the National Academy of Sciences. 2012;109:2418–2422.

18. Sweet SS, Pianka ER. Monitors, mammals and Wallace’s Line. Mertensiella. 2007;16:79–99.

19. Feit B, Gordon CE, Webb JK, Jessop TS, Laffan SW, Dempster T, Letnic M. Invasive cane toads might initiate cascades of direct and indirect effects in a terrestrial ecosystem. Biological Invasions. 2018;10.1007/s10550-018-1665-8.

20. Woinarski JCZ, Legge S, Fitzsimons JA, Traill BJ, Burbidge AA, Fisher A, et al. The disappearing mammal fauna of northern Australia: context, cause, and response. Conservation Letters. 2011;4:192–201.

21. Ziembicki MR, Woinarski JCZ, Webb JK, Vanderduys E, Tuft K, Smith J, et al. Stemming the tide: progress towards resolving the causes of decline and implementing management responses for the disappearing mammal fauna of northern Australia. THERYA. 2015;6:169–225.

22. McKenzie NL. Mammals of the Phanerozoic south-west Kimberley, Western Australia: biogeography and recent changes. Journal of Biogeography. 1981;8:263–280.

23. Braithwaite RW, Muller WJ. Rainfall, groundwater and refuges: predicting extinctions of Australian tropical mammal species. Austral Ecology. 1997;22:57–67.

24. Woinarski JCZ, Milne DJ, Wanganeen G. Changes in mammal populations in relatively intact landscapes of Kakadu National Park, Northern Territory, Australia. Austral Ecology. 2001;26:360–370.

25. McKenzie NL, Burbidge AA, Baynes A, Brereton RN, Dickman CR, Gordon G, et al. Analysis of factors implicated in the recent decline of Australia’s mammal fauna. Journal of Biogeography, 2007; 34:597–611.

26. Radford IJ, Dickman CR, Start AN, Palmer C, Carnes K, Everitt C, et al. Mammals of Australia’s tropical savannas: a conceptual model of assemblage structure and regulatory factors in the Kimberley region. PLOS ONE. 2014;9:0092341.

27. Frank A, Johnson CN, Potts JM, Fisher A, Lawes MJ, Woinarski JCZ, et al. Experimental evidence that feral cats cause local extirpation of small mammals in Australia’s tropical savannas. Journal of Applied Ecology, 2014;51:1486–1493.

28. McGregor HW, Legge S, Jones ME, Johnson CN. Landscape management of fire and grazing regimes alters the fine-scale habitat utilisation by feral cats. PLOS ONE. 2014;9:e109097.

29. McGregor HW, Legge S, Potts J, Jones ME, Johnson CN. Density and home range of feral cats in north-western Australia. Wildlife Research. 2015;42:223–231.

30. McGregor HW, Legge S, Jones ME, Johnson CN. Extraterritorial hunting expeditions to intense fire scars by feral cats. Scientific Reports. 2016;6:22559.

31. Leahy L, Legge SM, Tuft K, McGregor HW, Barmuta LA, Jones ME, Johnson, C. N. Amplified predation after fire suppresses rodent populations in Australia’s tropical savannas. Wildlife Research. 2016;42:705–716.

32. Dickman CR. Overview of the impacts of feral cats on Australian native fauna. Canberra: Australian Nature Conservation Agency. 1996;1–92.

33. Abbott I. Origin and spread of the cat, *Felis catus*, on mainland Australia, with a discussion of the magnitude of its early impact on native fauna. Wildlife Research. 2002;29:51–74.

34. Woinarski JCZ, Armstrong M, Brennan K, Fisher A, Griffiths AD, Hill B, et al. Monitoring indicates rapid and severe decline of native small mammals in Kakadu National Park, northern Australia. Wildlife Research. 2010;37;116–126.

35. Tuft K, Legge S, Frank ASK, James AI, May, T, Page E, Radford IJ, et al. Further experimental evidence that feral cats cause local extirpation of small mammals in Australia’s tropical savannas. Journal of Applied Ecology

36. Lawes MJ, Murphy BP, Fisher A, Woinarski JC, Edwards AC, Russell-Smith J. Small mammals decline with increasing fire extent in northern Australia: evidence from long-term monitoring in Kakadu National Park. International Journal of Wildland Fire. 2015;24:712–722.

37. Radford IJ, Gibson LA, Corey B, Carnes K, Fairman R. Influence of fire mosaics, habitat characteristics and cattle disturbance on mammals in fire-prone savanna landscapes of the northern Kimberley. PLOS ONE, 2015;10(6):p.e0130721.

38. Andersen AN, Cook GD, Corbett LK, Douglas MM, Eager RW, Russell-Smith J, et al. Fire frequency and biodiversity conservation in Australian tropical savannas: implications from the Kapalga fire experiment. Austral Ecology. 2005;30:155–167.

39. Shine R. The ecological impact of invasive cane toads (*Bufo marinus*) in Australia. The Quarterly Review of Biology. 2010;85:253–291.

40. Pearson DJ, Webb JK, Greenlees MJ, Phillips BL, Bedford GS, Brown GP, Thomas J, Shine R. Behavioural responses of reptile predators to invasive cane toads in tropical Australia. Austral Ecology, 2014;39:448–454.

41. Jolly CJ, Kelly E, Gillespie GR, Philips B, Webb JK. Out of the frying pan: Reintroduction of toad-smart northern quolls to southern Kakadu National Park. Austral Ecology. 2018;10.1111/aec.12551.

42. Brown GP, Phillips BL, Shine R. The ecological impact of invasive cane toads on tropical snakes: field data do not support laboratory-based predictions. Ecology. 2011;92:422–431.

43. Greenlees MJ, Brown GP, Webb JK, Phillips BL, Shine R. Effects of an invasive anuran [the cane toad (*Bufo marinus*)] on the invertebrate fauna of a tropical Australian floodplain. Animal Conservation. 2006;9:431–438.

44. Shine R, Koenig J. Snakes in the garden: an analysis of reptiles “rescued” by community-based wildlife carers. Biological Conservation. 200;102:271–283.

45. Cogger H. Reptiles and Amphibians of Australia. CSIRO Publishing. 2014.

46. Radford IJ, Fairman R. Fauna and vegetation responses to fire and invasion by toxic cane toads (*Rhinella marina*) in an obligate seeder dominated tropical savanna in the Kimberley, Northern Australia. Wildlife Research 2015;42:302–314.

47. Shine R. Ecology of eastern Australian whipsnakes of the genus *Demansia*. Journal of Herpetology. 1980;14:381–389.

48. Shine R, Slip D.J. Biological aspects of the adaptive radiation of Australasian pythons (Serpentes: Boidae). Herpetologica. 1990;46:283–290.

49. Shine R. Strangers in a strange land: ecology of the Australian colubrid snakes. Copeia. 1991;120–131.

50. Shine R. Allometric patterns in the ecology of Australian snakes. Copeia. 1994;851–867.

51. Shine R, Madsen T. Prey abundance and reproduction: rats and pythons on a tropical Australian floodplain. Ecology. 1997;78:1078–1086.

52. Zborowski P, Storey R. A Field Guide to Insects in Australia. Second Edition. Reed New Holland. 2003.

53. Liboschik T, Fried R, Fokianos K, Probst P. tscount: Analysis of Count Time Series. R package version 1.3.0. https://CRAN.R-project.org/package=tscount. 2016.

54. R Core Team. R: A language and environment for statistical computing. R Foundation for Statistical Computing, Vienna. https://www.R-project.org/. 2018.

55. Fokianos K, Fried R. Interventions in INGARCH processes. Journal of Time Series Analysis. 2010;31:210–225.

56. Ziembicki MR, Woinarski JC, Mackey B. Evaluating the status of species using Indigenous knowledge: Novel evidence for major native mammal declines in northern Australia. Biological Conservation. 2013;157:78–92.

57. Ibbett M, Woinarski JCZ, Oakwood M. Declines in the mammal assemblage of a rugged sandstone environment in Kakadu National Park, Northern Territory, Australia. Australian Mammalogy. 2017;doi:10.1071/AM17011.

58. Stokeld D, Fisher A, Gentles T, Hill B, Triggs B, Woinarski JCZ, Gillespie GR. What do predator diets tell us about mammal declines in Kakadu National Park? Wildlife Research. 2018;10.1071/WR17101.

59. Davies HF, McCarthy MA, Firth RSC, Woinarski JCZ, Gillespie GR, Andersen AN, et al. Declining populations in one of the last refuges for threatened mammal species in northern Australia. Austral Ecology. 2018;doi:10.1111/aec.12596.

60. Palmer C, Taylor R, Burbidge AA. Recovery Plan for the Golden Bandicoot, Isoodon auratus, and Golden-backed Tree-rat, Mesembriomys macrurus, 2004-2009. Department of Infrastructure, Planning & Environment; 2003.

61. Woinarski JC, Burbidge AA, Harrison PL. The action plan for Australian mammals 2012.

62. Burbidge AA, McKenzie NL. Patterns in the modern decline of Western Australia’s vertebrate fauna: causes and conservation implications. Biological conservation. 1989;50(1-4):143–98.

63. Woinarski JCZ, Murphy BP, Palmer R, Legge SM, Dickman CR, Doherty TS, et al. How many reptiles and killed by cats in Australia? Wildlife Research. 2018;45:247–266.

64. Legge S, Murphy BP, McGregor H, Woinarski JCZ, Augusteyn J, Ballard G, et al. Enumerating a continental-scale threat: How many feral cats are in Australia? Biological Conservation. 2017;206:293–303.

65. Woinarski JCZ, Murphy BP, Legge SM, Garnett ST, Lawes MJ, Comer S, et al. How many birds are killed by cats in Australia? Biological Conservation. 2017;214:76–87.

66. Woinarski JCZ, Legge SM, Dickman CR. Cats in Australia: companion and killer. CSIRO Publishing, Melbourne. 2019.

67. Slip DJ, Shine R. Habitat use, movements and activity patterns of free-ranging Diamond Pythons, *Morelia-spilota-spilota* (Serpentes, Boidae)-a radiotelemetric study. Wildlife Research. 1988;15(5):515–31.

68. Madsen T, Shine R. Seasonal migration of predators and Prey - A study of pythons and rats in tropical Australia. Ecology. 1996;77(1):149–56.

69. Brown GP, Shine R, Madsen T. Spatial ecology of slatey-grey snakes (*Stegonotus cucullatus*, Colubridae) on a tropical Australian floodplain. Journal of Tropical Ecology. 2005;21(6):605–12.

70. Corey B, Doody JS. Anthropogenic influences on the spatial ecology of a semi-arid python. Journal of Zoology. 2010;281:293–302.

71. Woinarski JCZ. The wildlife and vegetation of Purnululu (Bungle Bungle) National Park and adjacent area. Wildlife Research Bulletin. 1992(6).

72. Partridge TB. Fire and Fauna in Purnululu (Bungle Bungle) National Park, Kimberley, Western Australia. PhD Thesis. 2009.

73. Stokeld D, Gentles T, Young S, Hill B, Fisher A, Woinarski J, Gillespie G. Experimental evaluation of the role of feral cat predation in the decline of small mammals in Kakadu National Park. Final Report. Department of Environment and Natural Resources, Darwin. 2016.

74. Pianka ER. Diversity and adaptive radiations of Australian desert lizards. In Ecological Biogeography of Australia. Dr W. Junk Publishers, The Hague. 1981.

75. Morton SR, James CD. The diversity and abundance of lizards in arid Australia: a new hypothesis. The American Naturalist. 1988;132:237–256.

76. Gelman A. Scaling regression inputs by dividing by two standard deviations. Statistics in Medicine. 2008;27:2865–2873.

